# Biochemical and chemical biological approaches to mammalian sleep: roles of calcineurin in site-specific dephosphorylation and sleep regulation

**DOI:** 10.1101/2023.06.19.545643

**Authors:** Jianjun Yu, Tao V. Wang, Rui Gao, Chenggang Li, Huijie Liu, Lu Yang, Yuxiang Liu, Yunfeng Cui, Peng R. Chen, Yi Rao

## Abstract

Understanding of sleep mechanisms traditionally rely on electrophysiology and genetics but here we have initiated biochemical and chemical biological studies. Sleep was increased in mouse mutants with an alanine replacing threonine at residue 469 (T469A) of the salt inducible kinase 3 (SIK3). We searched for T469 phosphatases by classic purification with HEK293 cells and by a new photo-crosslinking method with mouse brains. Both led to PPP3CA, a catalytic subunit of calcium/calmodulin activated phosphatase (calcineurin). It dephosphorylated T469 and serine (S) 551 but not T221 in SIK3 in vitro. PPP3CA knockdown increased phosphorylation of T469 and S551 but not T221 in mouse brains. Knockdown of its regulatory subunit PPP3R1 significantly reduced daily sleep by more than 5 hours, exceeding other known mouse mutants. Our results have uncovered in vitro and in vivo evidence for site-specific SIK3 dephosphorylation by calcineurin, demonstrated a physiological role for calcineurin in sleep, and suggested sleep control by calcium dependent dephosphorylation.

---

Sleep is an important physiological process in animals^1^. It is regulated by the circadian and the homeostatic processes^2, 3^. The genetic approach has been taken in flies, mice, dogs and humans and proven powerful in uncovering many genes involved in sleep regulation^4–11^. Prominent examples include the discoveries of roles of orexin and its receptor in maintaining wakefulness^12,13^, and the finding of salt inducible kinase 3 (SIK3) through a forward genetic screen^14^. No attempt has been made to use biochemistry as a means of discovering molecules important for sleep because sleep could only be assayed in animals which are accessible to genetics, but not on molecules only biochemically accessible.

While we have taken the genetic approach in both Drosophila^15–18^ and the mouse^10, 17, 19^, we assessed its advantages and limits and decided to use both a biochemical approach with the classic purification method and a chemical biological approach with a newly developed chemical biological method to investigate molecular mechanisms of sleep by examining site-specific phosphorylation of a protein kinase previously found to be important for sleep regulation^14, 19–21^.

Protein phosphorylation differs between day and night and among different sleep-wake relevant states^22–25^. Multiple protein kinases in mammalian animals have been implicated in sleep regulation, including protein kinase A (PKA)^26–30^, the extracellular signal-regulated kinase (ERK)^31–33^, adenosine monophosphate (AMP)-activated protein kinase (AMPK)^34–36^, calcium (Ca^2+^)/calmodulin (CaM) kinase II (CaMKII) α and β^37–39^, c-Jun N-terminal kinase (JNK)^40^, SIK 3, 1 and 2^14, 19–21^, and the liver kinase B (LKB1)^17, 41–43^. A role for Ca^2+^-dependent pathway in regulating sleep has been proposed^38^ and was thought to be mediated at least in part by CaMKII α and β^38, 39^. The reduction of sleep per 24 hours (hrs) was more than 120 minutes (mins) in CaMK2β knockout mice^38^, which was more than other known genetic mutants in mice^10, 12–14, 17, 19, 21, 27, 30, 39, 44–53^.

While kinases are well known to play roles in mammalian sleep, little is known about protein phosphatases (PPases) in mammalian sleep. A protein discovered in the 1970s^54–56^ was named calcineurin (CaN, PP2 or PPP3)^57^ before its function as a phosphatase was characterized in the 1980s^58, 59^. By the 1990s, most of biochemical properties of CaN were known, such as that it is the only Ca^2+^- and calmodulin (CaM)-activated phosphatase, that the enzyme is a dimer made of a catalytic subunit calcineurin A (with three alternatives, PPP3CA, PPP3CB or PPP3CC) and a regulatory subunit calcineurin B (with two alternatives PPP3R1 or PPP3R2)^60–62^. PPP3CA, PPP3CB and PPP3R1 are ubiquitously expressed, whereas PPP3CC is abundant in the testis and PPP3R2 is specifically expressed in the testis. Loss of either PPP3CC or PPP3R2 leads to male infertility^63–68^. CaN functions in many systems, with its role in immunoregulation particularly striking^69–72^. Recently, the number of potential targets for all forms of calcineurin expanded from approximately 70 to 486 proteins^72^. In the brain, PPP3CA is the most abundant of the three catalytic subunits and PPP3R1 is the most abundant of the two regulatory subunits.

We began to search for PPases for SIK3 after defining the importance of a specific phosphorylation site in SIK3: threonine (T) 469. While loss of the serine (S) 551 equivalent sites in SIK1 and SIK2 caused the same gain of function (GOF) phenotype as that of S551 SIK3 GOF^21^, deletion of SIK1 or SIK2 gene did not affect sleep in mice^19^. We therefore have focused on SIK3 but not SIK1 or SIK2 in further studies. T469 and S551 of SIK3 are phosphorylated by PKA and their phosphorylation increased SIK3 interaction with the 14-3-3 protein^73^. 14-3-3 binding inhibited SIK3 and absence of either T469 or S551 increased SIK3 signaling^73^. S551 phosphorylation of SIK3 reduced sleep^14, 20^. The functional significance of T469 phosphorylation in SIK3 was previously unknown. Here we have generated T to alanine (A) mutation of the amino acid residue at 469 (T469A) of SIK3. We found sleep increased in T469A mice as compared to wild type (wt) mice, indicating that the function of T469 is similar to that of S551 in sleep regulation.

We used two approaches to search for PPases of T469 and S551 of SIK3: classic biochemistry of purifying PPases for T469 and S551 of SIK3 from human embryonic kidney (HEK) HEK 293T cells, and chemical biology of photo-crosslinking of proteins (RG and PRC) interacting with SIK3 in the mouse brain. In both experiments, PPP3CA, a catalytic subunit of CaN, was found. in vitro biochemical assays demonstrated that PPP3CA dephosphorylated phospho-T469 and phospho-S551 of SIK3. This reaction also required Ca^2+^, CaM and PPP3R1. Interestingly, PPP3CA did not dephosphorylate T221 of SIK3, a site whose phosphorylation promotes SIK3 activity^17, 19^ and indicates sleep need and promotes sleep^19^. Ca^2+^ ionophore induced dephosphorylation of T469 and S551, but not T221, was inhibited in HEK cells when PPP3CA and PPP3CB or the regulatory subunit PPP3R1 was knocked down by small guided RNA (sgRNA) mediated gene knockdown. Phosphorylation of T469 and S551, but not T221, was increased in mouse brains when PPP3CA or PPP3R1 was knocked down, indicating that PPP3CA and PPP3R1 is physiologically required for dephosphorylation of T469 and S551 in vivo. Knockdown of PPP3CA reduced sleep by approximately 3 hrs (or 187.6±13.0 mins to be precise) over 24 hrs. Knockdown of PPP3R1 reduced sleep by more than 5 hrs (or 349.3±21.5 mins) per day, which exceeded sleep changes in all mouse mutants tested so far. Measured by the extent of changes in sleep, CaN is the most significant regulator of sleep identified so far. Our work is the first to rely on biochemistry and chemical biology to discover molecules important for sleep and it has shown the awesome power of biochemistry and chemical biology (in addition to that of genetics) in sleep research.

## Results

### Functional significance of T469 in regulating mouse sleep

To study the potential role of T469 in SIK3, we generated a line of mice carrying the T469A mutation (Fig. 1, Extended Data Figs. 1 and 2, Extended Data Tables 1 and 2). Sleep is usually measured in male mice but, because male mice with the genotype of SIK3^T469A/T469A^ were embryonic lethal, we could only compare the sleep phenotypes of two genotypes (SIK3^+/+^ and SIK3^T469A/+^) in males. We have also examined the sleep phenotypes of all three genotypes (SIK3^+/+^ and SIK3^T469A/+^ and SIK3^T469A/T469A^) in females (Extended Data Figs. 3 and 4, Extended Data Tables 1 and 2). The phenotypes in males and females were qualitatively similar, though the phenotypes of SIK3^T469A/T469A^ were stronger than those of SIK3^T469A/+^, as expected.

**Fig. 1.**
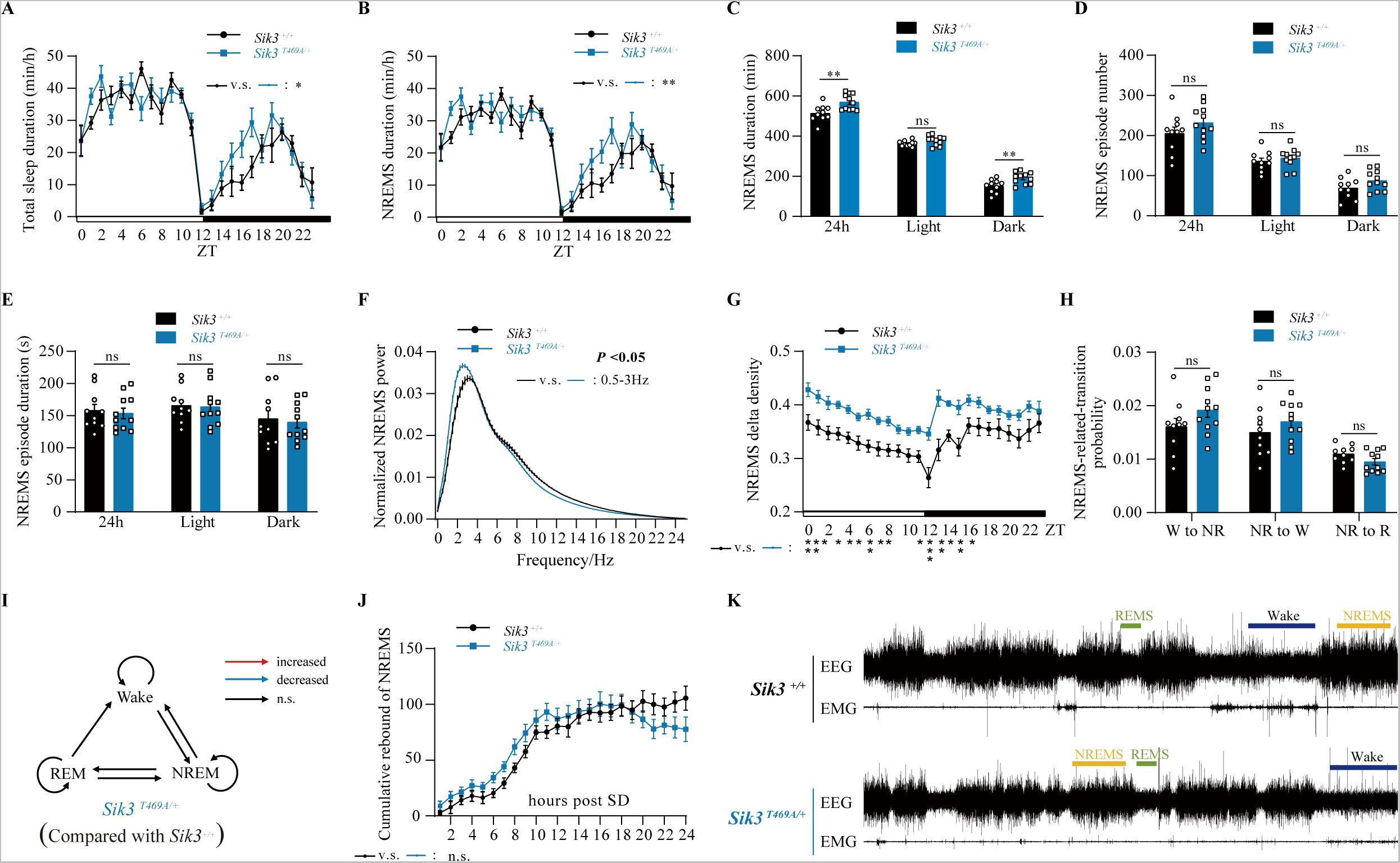
Sleep phenotype of T469A mutant male mice. Sleep phenotype of male mice with either SIK3^+/+^ genotype or SIK3^T469A/+^ genotype. SIK3^T469A/ T469A^ males were embryonic lethal although females of all three genotypes including SIK3^T469A/ T469A^ were viable and analyzed in Extended Data Figs. 3 and 4. For detailed numbers, see Extended Data Tables 1 and 2. **a, b**, profiles showing sleep time each hour in min/hr (**a**) or profiles of NREM sleep (**b**). The x axis shows zeitgeber time (ZT) with the white box indicating light phase (or daytime) and black box dark phase (or nighttime). The black line shows data from SIK3^+/+^ mice (n = 11), the blue line data from SIK3^T469A/+^ mice (n = 10). ns: statistically not significant. *p<0.05; **p<0.01; mean ± standard error of the mean (SEM) (Two-way ANOVA with Tukey’s multiple comparisons test). **c,d,** Data and statistics of NREM sleep duration (**c,** *p< 0.05, **p<0.01; ns, not significant; mean ± SEM, One-way ANOVA with Tukey’s multiple comparisons test), NREM sleep episode number (**d**, ns, not significant; mean ± SEM, Kruskal-Wallis test with Dunn’s multiple comparisons test), and NREM sleep episode duration (**e**, ns, not significant; mean ± SEM, Kruskal-Wallis test with Dunn’s multiple comparisons test). **f**, EEG power spectrum during NREM sleep. X-axis indicates frequency distribution of EEG power. *p<0.05; mean ± SEM (Two-way repeated measurement ANOVA with Tukey’s multiple comparisons test). **g**, Diurnal NREMS delta density. *p<0.05; **p<0.01; ***p<0.001; ns, not significant; mean ± SEM (Mixed-effects model) NREMS delta power density over 24 hours. X-axis indicates ZT. **h**, Probabilities of transition between different sleep and wake states. ns, not significant; mean ± SEM (Two-way ANOVA with Tukey’s multiple comparisons test). **i**, Recovery of NREM sleep after 6 hrs of SD. ns, not significant; mean±SEM (Two-way repeated measurement ANOVA with Tukey’s multiple comparisons test). **j**, One hr representative EEG and EMG graphs at different vigilance states (wake, NREM, REM).

Representative electroencephalogram (EEG) and electromyogram (EMG) graphs are shown in Fig. 1K and typical hypnograms shown in Figure S2E. Over 24 hrs, SIK3^T469A/+^ males slept more than SIK3^+/+^ males by 50.1±18.9 mins (Fig. 1a, Extended Data Table 2). During daytime, (or the light phase), there was no significant difference in sleep time between SIK3^T469A/+^ and SIK3^+/+^ males (Fig. 1a, Extended Data Table 2, 7.7±9.7 mins). During nighttime (or the dark phase), SIK3^T469A/+^ males slept more than SIK3^+/+^ males by 42.4±15.1 mins (Fig. 1a, Extended Data Table S). The amount of total non-rapid eye movement (NREM) sleep during 24 hrs was increased in SIK3^T469A/+^ males by 55.8±17.5 mins over SIK3^+/+^ males (Fig. 1 b and c). The amount of nighttime NREM sleep was increased in SIK3^T469A/+^ males by 40.6±13.6 mins over SIK3^+/+^ males (Fig. 1 b and c). Neither NREM episode number nor episode duration reached statistical significance (Fig. 1 d and e). Rapid eye movement (REM) sleep was not significantly different between SIK3^T469A/+^ and SIK3^+/+^ males (Extended Data Fig. 1 e-g, Extended Data Table 2). Power spectrum of EEG showed only increase in the 0.5 to 3 Hertz (Hz) range (Fig. 1f). Sleep need is an important regulator of sleep^2, 3, 74, 75^. It is regulated by prior wakeful experience^2, 76^ and can be measured by NREM delta power densities which is a measure of EEG activity in the 1-4 Hz range^2, 74, 75, 77, 78^. NREM delta power densities are significantly increased in SIK3^T469A/+^ males (Fig. 1g). Other parameters were not significantly different between SIK3^T469A/+^ and SIK3^+/+^ males.

In females, SIK3^T469A/T469A^ mice showed significantly more NREM sleep in the dark phase (84.6±26.5 mins more than SIK3^+/+^ mice or 60.8±27.4 mins more than SIK3^T469A/+^ mice) (Extended Data Fig. 3 a and b, Extended Data Tables 1 and 2), attributable to an increase of NREM episode number (Extended Data Fig. 3c) but not episode duration (Extended Data Fig. 3d). REM sleep was not significantly different in the dark phase, but decreased in the light phase (Extended Data Fig. 3 e and f, Extended Data Tables 1 and 2), attributable to the decrease of REM episode number (Extended Data Fig. 3g) but not episode duration (Extended Data Fig. 3h). Wake was decreased in the dark but not the light phase (Extended Data Fig. 3 i and j, Extended Data Tables 1 and 2). Wake episode number was increased in the dark (Extended Data Fig. 3) while wake episode duration was decreased in the dark (Extended Data Fig. 3l). Transition from NREM to REM was decreased whereas the other transitions were not different (Extended Data Fig. 3 m to p). NREM delta power densities were increased in both SIK3^T469A/T469A^ and SIK3^T469A/+^ females when compared to SIK3^+/+^ females (Extended Data Fig. 4).

In summary, in male and female T469A mutants, both total sleep and NREM sleep in the dark phase was increased. NREM delta power densities were also increased in both male and female T469A mutant mice. T469 phosphorylation is indeed a negative regulator of basal NREM sleep and sleep need, which are both opposite to the roles of SIK3^14, 19–21^.

### Biochemical purification of phosphatase(s) for T469 and S551 of SIK3

With HEK 293T cells, we biochemically purified phosphatase(s) capable of dephosphorylating T469 and S551 of SIK3 (Fig. 2).

**Fig. 2.**
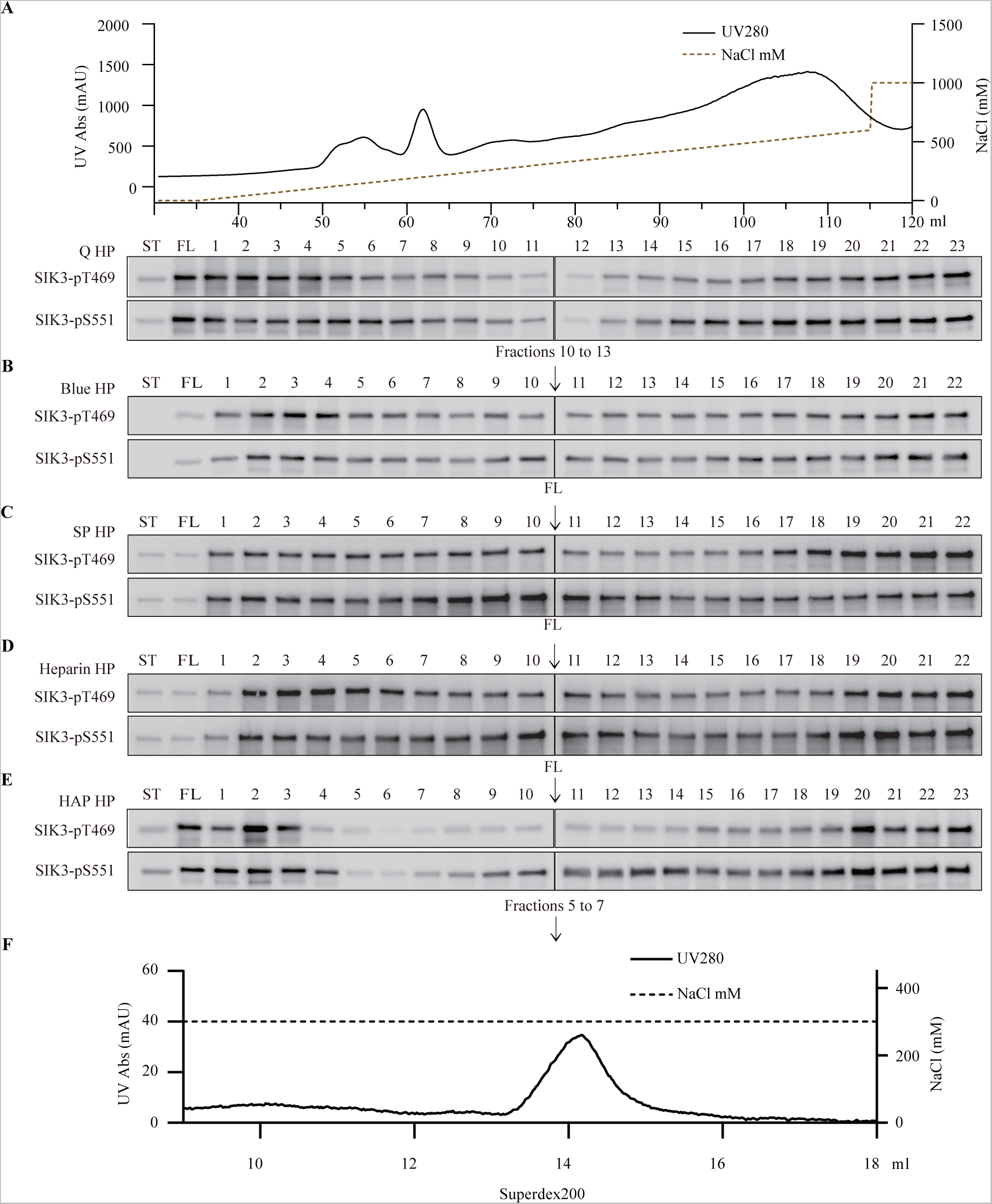
Biochemical purification of SIK3 phosphatases from HEK cells. **a**, 500 mg (at a concentration of 10 mg/ml) of HEK cell lysates was fractionated on a QHP column, eluted with a linear gradient of NaCl (0-600 mM) and the final wash with 1 M NaCl buffer A. Each fraction was dialyzed into buffer A, removing NaCl. 10 μl of each sample was used for analysis of activities removing phosphate from SIK3 T469 and S551. T469 and S551 of bacterially expressed recombinant SIK3 were phosphorylated by PKA in vitro before being used to test phosphatase activities of the fractions of HEK lysates. Fractions 10 to 13 contained significant activities removing phosphate from SIK3 T469 and S551. **b**, Fractions 10 to 13 from QHP were combined and dialyzed with buffer A before being loaded onto a Blue HP column. It was eluted with a linear gradient of NaCl (0-600 mM) and the final wash with 1 M NaCl buffer A. The flow through (FL) fraction from the Blue HP contained significant activities removing phosphate from SIK3 T469 and S551. **c,** The FL fraction from the Blue HP column was dialyzed with buffer A and loaded onto a SP-HP column. The fractionation was similar to those in (**a**) and (**b**). The FL fraction from the SP-HP column contained significant activities removing phosphate from SIK3 T469 and S551. **d,** The FL fraction from the SP-HP column was dialyzed with buffer A and loaded onto a heparin column. The rest of the fractionation was similar to (**a**) and (**b**). The FL fraction and fraction 1 contained significant activities removing phosphate from SIK3 T469 and S551. **e,** The FL fraction from the heparin column was dialyzed with buffer A and loaded onto a HAP-HP column. The rest of the fractionation was similar to (**a**) and (**b**) except that the final wash was with 5 CVs of 500 mM K_2_PO_4_. Fractions 5 to 7 contained significant activities removing phosphate from SIK3 T469 and S551. **f,** Active fractions from the HAP-HP column were condensed into 0.5 ml, fractionated on a Superdex 200 molecular sieve column, eluted with 300 mM NaCl gradient into 20 CVs. 1 ml from each fraction was collected and labeled as samples 1 to 20. Protein contents were monitored with UV at 280 nm.

We first ensured that the antibodies we used could specifically recognize the phospho-forms of T469 and S551 of SIK3. Recombinant SIK3 fragment containing its amino acid residues (aa) 1 to 558 was generated in and purified from *Escherichia coli* (*E. coli*). The presence of both PKA and ATP catalyzed phosphorylation of T469 and S551 in vitro. When the antibodies were tested, they were each confirmed to be specific for the phosphorylated forms of T469 and S551, respectively (Extended Data Fig. 5).

We lysed HEK 293T cells and fractionated 500 mg of cell lysates (at a concentration of 10 mg/ml) sequentially on a Q HP, a Blue HP, an SP HP, a heparin HP, a hydroxyapatite (HAP) HP and a Superdex 200 column (Fig. 2 and Fig. 3a). At each step, an aliquot from each fraction was assayed for PPase activities at T469 and S551. Active fractions from one step were combined and further fractionated on the next column. Thus, fractions 10 to 13 from the Q HP column, the flow through (FL) fraction from the Blue HP column, the FL from the SP HP column, the FL from the heparin column, and fractions 5 to 7 from the HAP HP column were each loaded onto the next column (Fig. 2).

**Fig. 3.**
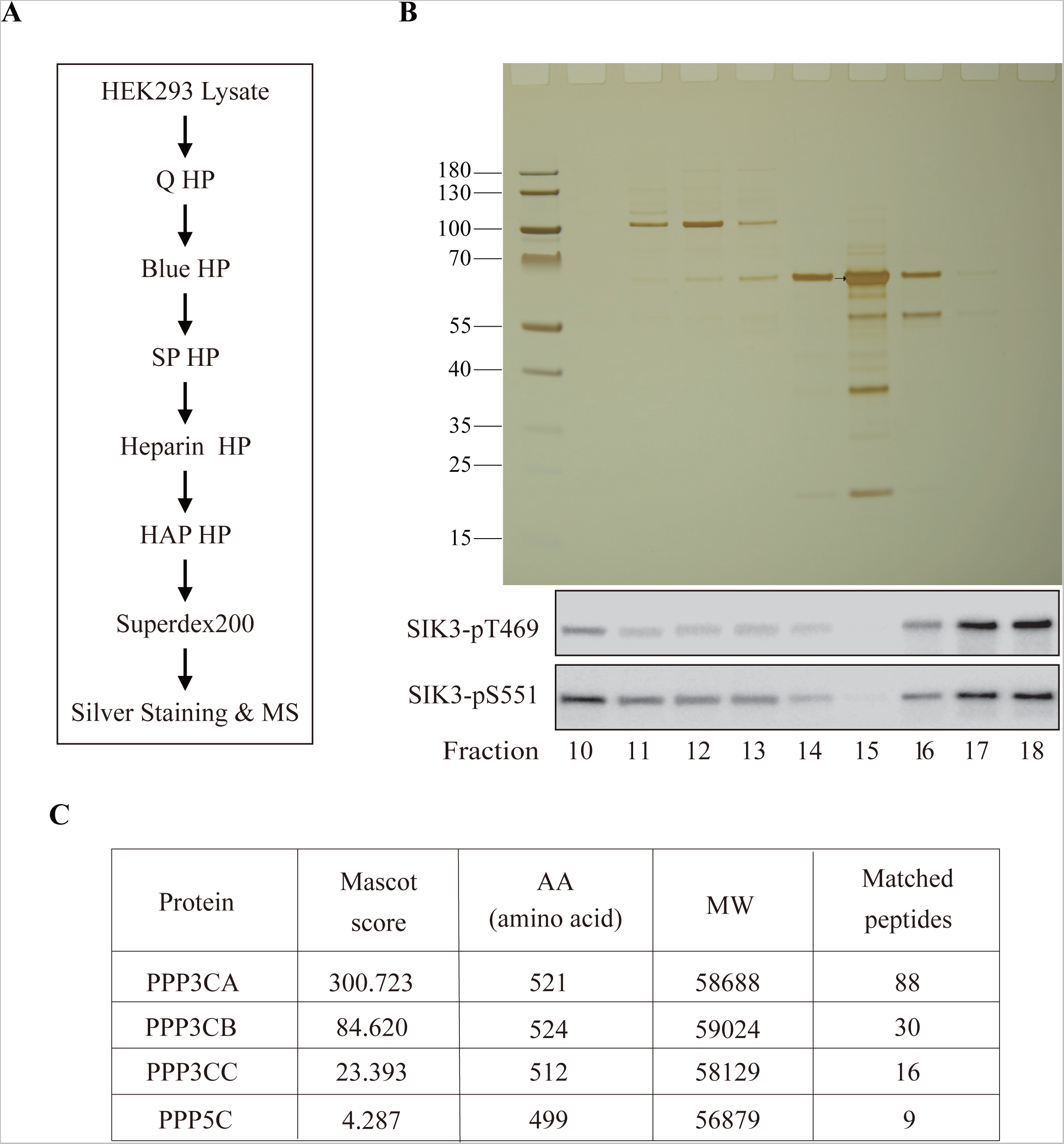
Identification of protein phosphatases purified from HEK cells. **a,** A schematic illustration of phosphatase purification with HEK cell lysates passing through Q HP, Blue HP, SP HP, heparin HP, HAP HP and Superdex 200 columns before silver staining and mass spectroscopic analysis. **b,** Fractions from the Superdex 200 column with strongest phosphatase activities for T469 and S551 in Fraction 15. 25 μl of each fraction (from Fraction 10 to Fraction 18) was run onto a gel and silver-stained. The thick band indicated by the arrow was analyzed by mass spectroscopy. **c,** 4 protein phosphatases were found by MS.

During purification, we observed that PPase activities for T469 were highly correlated with those for S551 (Fig. 2 and Extended Data Fig. 3b).

Fractions 10 to 18 from the Superdex 200 column were run onto a polyacrylamide gel and silver-stained for proteins (Fig. 3b). Fraction 15 contained the strongest PPase activities. A thick band indicated by an arrow in the lane for fraction 15 was cut and sent for mass spectroscopic (MS) analysis. Four PPases were detected: PPP3CA, PPP3CB, PPP3CC and PPP5C (Fig. 3c).

### Discovery of PPP3CA as a protein-protein interaction partner for SIK3 in the mouse brain

We used a new photo-crosslinking method invented by two of us (RG and PRC, unpublished) to search for proteins interacting with SIK3 in vivo. We intended to capture interacting proteins of SIK3 in the brain. Compared with digested neuronal cell culture or cell lysates, brain slices are better in keeping in situ protein networks. Among current approaches for studying protein-protein interactions (PPIs), photo-crosslinking strategies which covalently capture interacting proteins under light irradiation are considered to be the more desired methods in living systems due to their good temporal resolution and low cytotoxicity compared with chemical crosslinking. However, the incorporation of photo-crosslinkers into proteins of interests (POIs) normally relies on genetic code expansion (GCE) strategy to engineer native tRNA synthetases for photoactivable unnatural amino acid insertion^79^ or metabolic turn-over processes for photoactivable moiety containing amino acid analogues^80^, which are both difficult to achieve in tissue samples. Furthermore, traditional moieties used for photo-crosslinking such as diazirine and aryl azide are normally sensitive to ultraviolet (UV) irradiation^81^, which possesses high-energy phototoxicity and weak tissue penetration^82^. As a result, photo-crosslinking strategy for in situ PPI capture in tissue level samples has not been frequently reported.

Based on the previous work from the Chen lab^83^, here we (RG and PRC) have developed a photocatalytic crosslinking strategy for capturing PPIs in mouse brain slices (Fig. 4a). Considering the challenges as mentioned, we chose eosin Y as the photocatalyst and 1,6-diaminohexane as the linker. Each compound shows good solubility and eosin Y, in particular, is widely used in tissue staining. With maximum absorption at 517 nm, eosin Y generates singlet oxygen (^1^O_2_) under green light irradiation^84^, which activates side chains of certain amino acid residues such as tyrosine to form electrophilic intermediates which could further form covalent linkage with amine warhead^85^. Our method does not require transfection or genetic modification to incorporate the photocatalyst or the crosslinker into target samples for PPI capture.

**Fig. 4.**
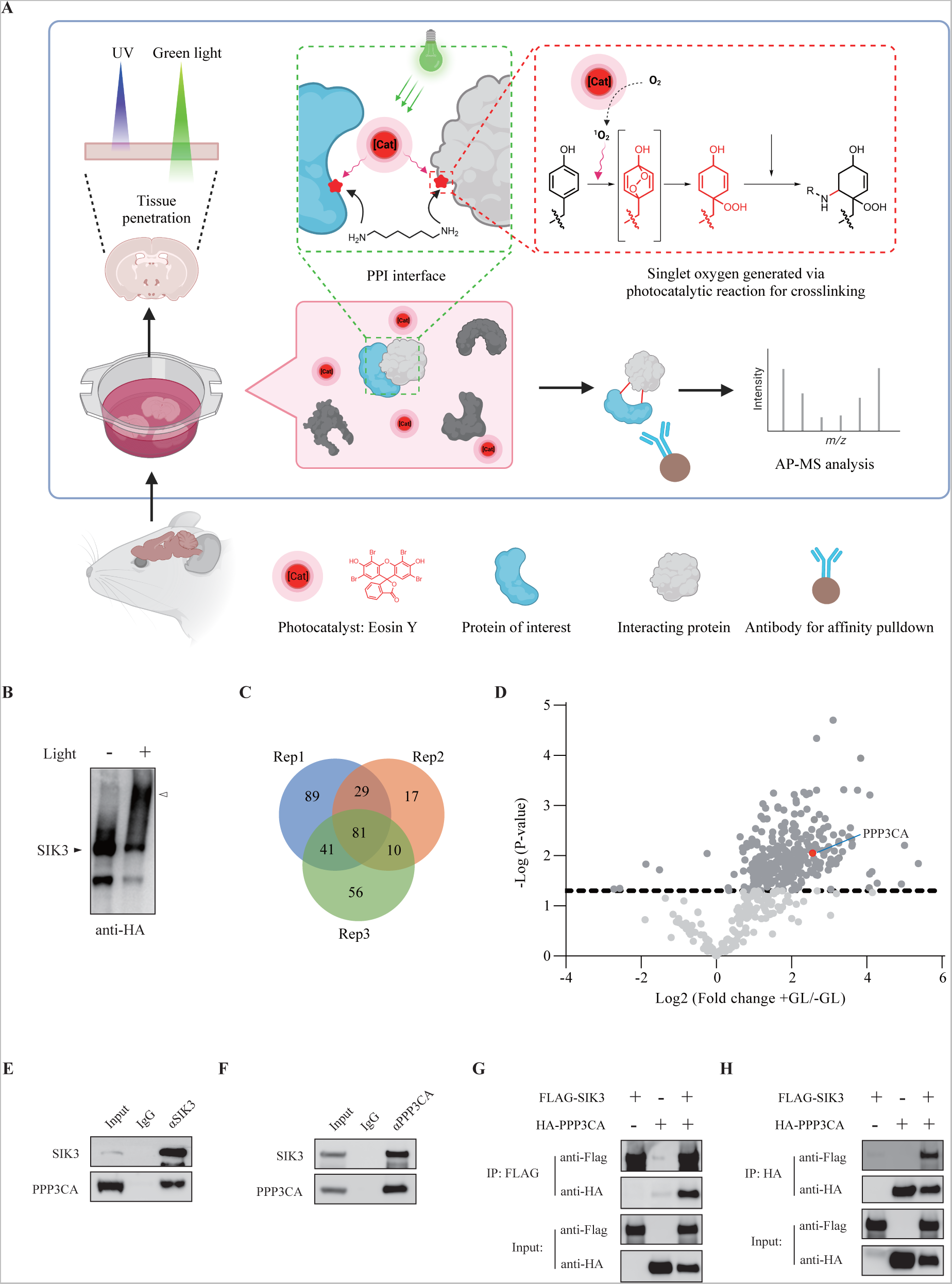
Proteins interacting with SIK3 identified by photo-crosslinking. **a,** A schematic diagram of our newly invented photo crosslinking method (RG and PC) (for details, see the method section). We have generated a mouse line with the SIK3 tagged at its C terminus with 3 repeats of the HA epitope. Brain slices were prepared from SIK3-HA mice and photocatalytic crosslinking was carried out in darkness with eosin Y (50 μM) and 1, 6 dihexamine (1 mM). The anti-HA antibody was used to pull down proteins crosslinked to SIK3. **b,** Comparison of proteins pulled down by anti-HA before and after light treatment. Validation of photocatalytic crosslinking efficacy in mouse brain slices. 1 hr green light was given after photocatalytic reagents treatment and slices were collected and homogenized. The arrowhead indicates SIK3-interaction proteins. **c**-**d**, Veen (**c**) and volcano (**d**) plot of putative SIK3 interaction proteins enriched in photo-linkage groups. The red dot is PPP3CA. **e**-**f**, Co-immunoprecipitation between PPP3CA and SIK3 from WT mouse brain homogenates using anti-SIK3 and anti-PPP3CA antibodies. **g**-**h,** Co-immunoprecipitation between the HA tagged PPP3CA and FLAG tagged SIK3 in HEK293T cells using anti-FLAG and anti-HA antibodies. HEK293T cells were co-overexpressed with 1 μg HA-PPP3CA and 1 μg FLAG-SIK3 expressing plasmids for 24 hrs before cell collection and lysis.

We have generated SIK3-3xHA mice in which the SIK3 protein was tagged with a hemagglutinin (HA) epitope at its carboxyl (C) terminus^86^. This allowed us to examine proteins associated with SIK3-HA after photo-crosslinking (Fig. 4b).

We applied the photocatalytic crosslinking strategy in freshly prepared mouse brain slice samples of the SIK3-HA mice. Eosin and 1,6-diaminohexane were incorporated into samples following the protocol described, followed by green LED (GL) irradiation. The SIK3 interactome was verified by Western analysis, of which clear crosslinking bands were presented in the +GL group, indicating efficient activation of our probes (Fig. 4b).

For both crosslinking and non-crosslinking groups, SIK3 protein was enriched by anti-HA beads and the crosslinking interactome could be pulled down at the same time. For further analysis of the proteins in crosslinking complexes, we excised bands with molecular weight larger than SIK3 for MS sample preparation. Quantitative studies were performed by applying dimethyl labeling method towards both groups after trypsin digestion. Although there were individual variations, 81 proteins overlapped in three independent experiment replications with high level of enrichment in the +GL groups (ratio +GL/-GL>4, log2(+GL/-GL)>2), just as shown in the Venn diagram (Fig. 4c). To better justify the protein candidates, we applied a volcano plot analysis (Fig. 4d), in which only the proteins with fold-change more than 4 and p-value less than 0.05 were considered as significant interacting candidates of SIK3. There was only one catalytic PPase: PPP3CA (Fig. 4d).

To confirm interaction of SIK3 and PPP3CA in the brain, we used antibodies for SIK3 in an immunoprecipitation experiment and found both SIK3 and PPP3CA in the precipitates (Fig. 4e). Similarly, both PPP3CA and SIK3 were found in the precipitates when anti-PPP3CA antibodies were used to immunoprecipitate brain lysates (Fig. 4f). This interaction could be recapitulated in HEK 293T cells when FLAG tagged SIK3 and HA tagged PPP3CA were introduced into HEK cells and immunoprecipitation experiments carried out with either the anti-FLAG (Fig. 4g) or the anti-HA (Fig. 4h) antibodies.

### in vitro Dephosphorylation of T469 and S551, but not T221 by PPP3CA

While the biochemical approach has uncovered 4 PPases with potential activities on SIK3, and the chemical biological approach has uncovered PPP3CA as an interacting protein for SIK3, it was unclear which PPase(s) could dephosphorylate SIK3 at T469 or S551. Given the widely known assumption that PPases are notoriously “non-specific”, it was also unclear whether any of the PPases identified by us was specific.

We immunoprecipitated FLAG tagged SIK3 expressed in HEK 293T cells as the substrate for testing PPases because it was phosphorylated at T221, T469 and S551 (Fig. 5a). Each of the PPases (PPP3CA, PPP3CB, PPP3CC, PPP5C and PPP3R1) was also expressed in and immunoprecipitated from HEK cells. T469 and S551 of SIK3 could be dephosphorylated by PPP3CA, PPP3CB and PPP3CC, in the presence of PPP3R1. By contrast, PPP3CA, PPP3CB and PPP3CC could not dephosphorylate T221 of SIK3. Under the same conditions, PPP5C could not dephosphorylate T469, S551 or T221 of SIK3 (Fig. 5a).

**Fig. 5.**
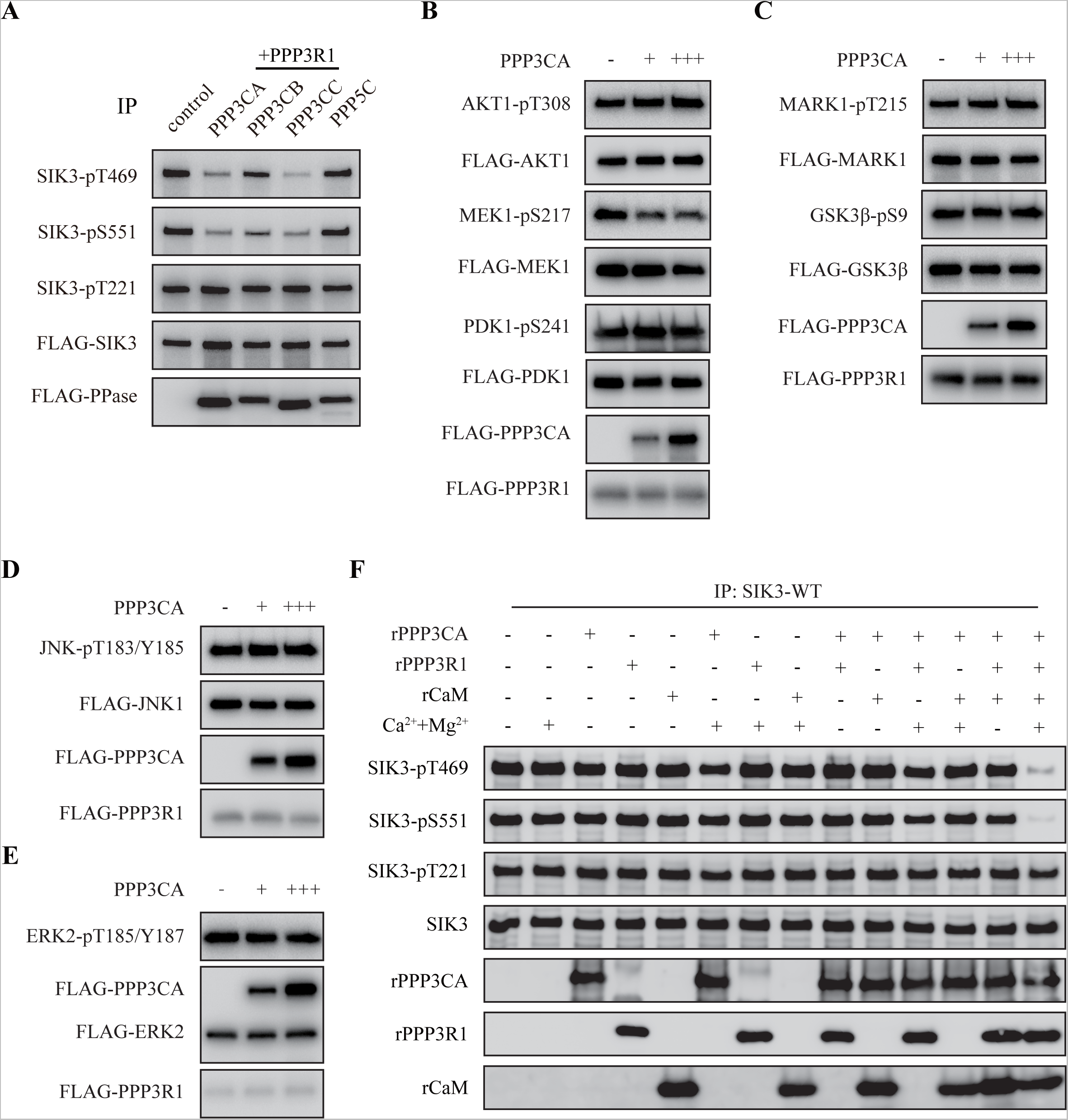
Site-specific dephosphorylation of SIK3 by PPP3CA in vitro. **a,** PPP3CA, PPP3CB, PPP3CC and PPP5C were individually tagged with FLAG and transfected into HEK 293T cells. PPP3R1 was also separately transfected into HEK 293T cells. SIK3-FLAG was transfected into HEK 293T cells. Each of the PPases and the regulatory PPP3R1 was separately immunoprecipitated by the anti-FLAG antibody from HEK cells. Under the same conditions (in the presence of PPP3R1), PPP3CA, PPP3CB or PPP3CC could dephosphorylate SIK3 at T469 and S551, but not T221. PPP5C could no dephosphorylate SIK3 at T469, S551or T221.**b-e,** In the presence of PPP3R1 immunoprecipitated from HEK cells, increasing concentrations (from + to +++) of PPP3CA immunoprecipitated from HEK cells could not dephosphorylate AKT1 at T308, PDK1 at S241, MARK1 at T215, GSK3β at S9, JNK1 at T183, ERK2 at T185. In the presence of PPP3R1 immunoprecipitated from HEK cells, PPP3CA immunoprecipitated from HEK cells could dephosphorylate MEK1 at S217 (**b**). **f,** Recombinant PPP3CA, PPP3R1, CaM were expressed in and purified from *E. coli*. Full-length SIK3 was immunoprecipitated from HEK 293T cells. In the presence of Ca^2+^, PPP3R1 and CaM, PPP3CA dephosphorylated SIK3 at T469 or S551but not at T221.

Because PPP3CA and PPP3R1 are abundant in the brain while PPP3CC and PPP3R2 are thought to be enriched or specific in the testis^63–68^, we focused on PPP3CA and PPP3R1. In the presence of PPP3R1, PPP3CA immunoprecipitated from HEK cells could dephosphorylate MEK1 at S217, but not T308 of AKT1, S241 of PDK1, T215 of MARK1, S9 of GSK3β, T183 of JNK1, T185 of ERK2 (Fig. 5 b-e).

Experiments in Fig. 5 a to e used enzymes immunoprecipitated from HEK cells and it could not be ruled out that they contained other proteins associated with the intended PPase. We expressed CaM, PPP3CA and PPP3R1 in *E. coli* so that the absence of protein serine/threonine PPases in *E. coli* made the contamination impossible. We found that CaM, PPP3CA, PPP3R1 and Ca^2+^ were all required for dephosphorylation of T469 and S551 of SIK3 (Fig. 5f). This further showed that CaN is a Ca^2+^/CaM-activated PPase requiring both its catalytic PPP3CA subunit and regulatory PPP3R1 subunit.

Thus, our in vitro results have demonstrated that PPP3CA is highly specific, because not only it does not dephosphorylate tested kinases other than SIK3 and MEK1, but it also targets T469 and S551 but not T221 in SIK3.

### in vivo Requirement of PPP3CA and PPP3R1 for dephosphorylation of T469 and S551, but not T221 in HEK cells

We investigated whether PPP3CA could dephosphorylate T469 and S551 of SIK3 in HEK293T cells.

Ionomycin is a classic Ca^2+^ ionophore, allowing Ca^2+^ flow into cells. When applied to media culturing HEK cells, ionomycin could dephosphorylate T469 and S551 of SIK3 in a dose- (Fig. 6a) and time- (Fig. 6b) sensitive manner. It was also specific without evidence for changes in SIK3-T221 phosphorylation (Fig. 6 a-b).

**Fig. 6.**
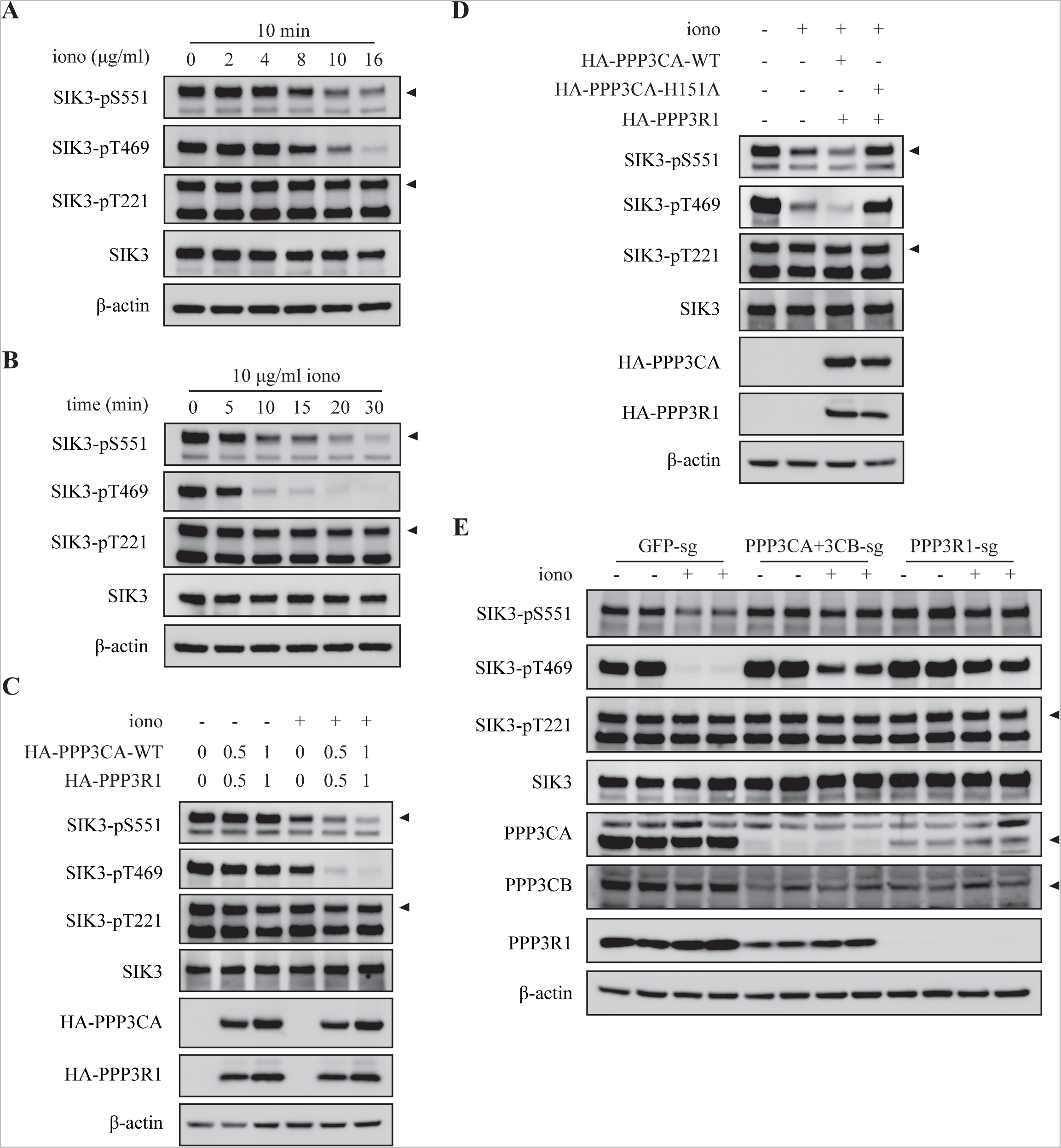
>SIK3 dephosphorylation and PPP3CA in HEK cells. **a-b,** Ionomycin dephosphorylated SIK3 at T469 and S551 (but not T221) in a dose-(**a**) and time- (**b**) dependent manner. **c**, Transfection of increasing dosage of PPP3CA and PPP3R1 into HEK cells enhanced the dephosphorylation response of SIK3-T469 and SIK3-S551 to ionomycin. In the absence of ionomycin, increasing the dosages of PPP3CA and PPP3R1 alone did not significantly changed SIK3-T469 and SIK3-S551 phosphorylation. After transfection for 24 hrs, cells were treated with 10 μg/ml ionomycin for 10 min. **d**, Transfection of a catalytically inactive PPP3CA mutant (H151A) and PPP3R1 into HEK cells did not enhance the dephosphorylation response of SIK3-T469 and SIK3-S551 to ionomycin. **e,** Transfection of sgRNA targeting GFP into HEK 293T cells did not significantly affect the dephosphorylation response of SIK3-T469 and SIK3-S551 to ionomycin. Transfection of sgRNA targeting either PPP3CA or PPP3CB alone into HEK 293T cells affected the dephosphorylation response of SIK3-T469 and SIK3-S551 to ionomycin in a variable manner (data not shown). Transfection of sgRNAs targeting both PPP3CA and PPP3CB into HEK 293T cells robustly and significantly reduced the dephosphorylation response of SIK3-T469 and SIK3-S551 to ionomycin. Transfection of sgRNA targeting PPP3R1 into HEK 293T cells robustly and significantly reduced the dephosphorylation response of SIK3-T469 and SIK3-S551 to ionomycin. PPP3R1 targeting reduced the protein levels of itself as well as PPP3CA, and PPP3CB in HEK cells. PPP3CA and PPP3CB targeting reduced the protein levels of themselves as well as PPP3R1 in HEK cells, indicating the inter-dependence of CaN catalytic and regulatory subunits.

Overexpression of PPP3CA/PPP3R1 enhanced the ionomycin induced dephosphorylation of T469 and S551 in a dosage-dependent manner (Fig. 6c). However, overexpression of an inactive form of PPP3CA, PPP3CA-H151A (with histidine, or H, mutated to A), did not dephosphorylate T469 and S551 (Fig. 6d).

To investigate calcineurin dependence of ionomycin induced dephosphorylation, we used sgRNAs to generate single or double knockout HEK293T cell lines for PPP3CA, PPP3CB and PPP3R1. The phenotype of single knockout for PPP3CA or PPP3CB in HEK cells was variable. However, phenotypes of PPP3CA and PPP3CB double knockout, or PPP3R1 single knockout, in HEK cells was robust. The ionomycin induced dephosphorylation of T469 and S551 of SIK3 was significantly inhibited by PPP3R1 knockout or PPP3CA and PPP3CB double knockout in HEK cells (Fig. 6e). Again, the effect was specific as phosphorylation of T221 of SIK3 was not significantly affected in any of these knockout HEK cells.

### in vivo Requirement of PPP3CA and PPP3R1 for dephosphorylation of T469 and S551, but not T221, in the mouse brain

To investigate their in vivo roles in the mouse, we used the CRISPR-Cas9 strategy to target PPP3CA or PPP3R1 in the brains of adult mice, with three sgRNAs for each gene (Extended Data Fig. 6). There were two types of controls: WT^Ctrl^ were wt mice injected with the same viruses as those for targeting PPP3CA or PPP3R1; eGFP^Ctrl^ were mice with Cas9 expressed from the Rosa26 locus (Rosa^Cas9/+^) injected with sgRNAs targeting the enhanced green fluorescent protein (eGFP). PPP3CA knockdown (PPP3CA^KD^) mice were generated by injecting Rosa^Cas9/+^ mice with AAV2/PHP.eB-CMV-mScarlet-PPP3CA-sgRNA-WPRE. PPP3R1^KD^ mice were generated by injecting Rosa^Cas9/+^ mice with AAV2/PHP.eB-CMV-mScarlet-PPP3R1-sgRNA-WPRE.

Knockdown efficiency was examined by Western analysis, which showed that sgRNAs targeting PPP3CA or PPP3R1 readily decreased their protein levels. Furthermore, when PPP3CA was targeted by sgRNA, levels of PPP3CA and PPP3R1 proteins were reduced (Fig. 7a), showing interdependence of the catalytic and the regulatory subunits. However, PPP3CB protein level was not reduced in PPP3CA^KD^ mice (Fig. 7a), indicating the specificity of sgRNAs for PPP3CA over PPP3CB. In PPP3R1^KD^ mice, levels of PPP3R1, PPP3CA and PPP3CB were reduced (Fig. 7b), again confirming the interdependence of catalytic and regulatory subunits.

**Fig. 7.**
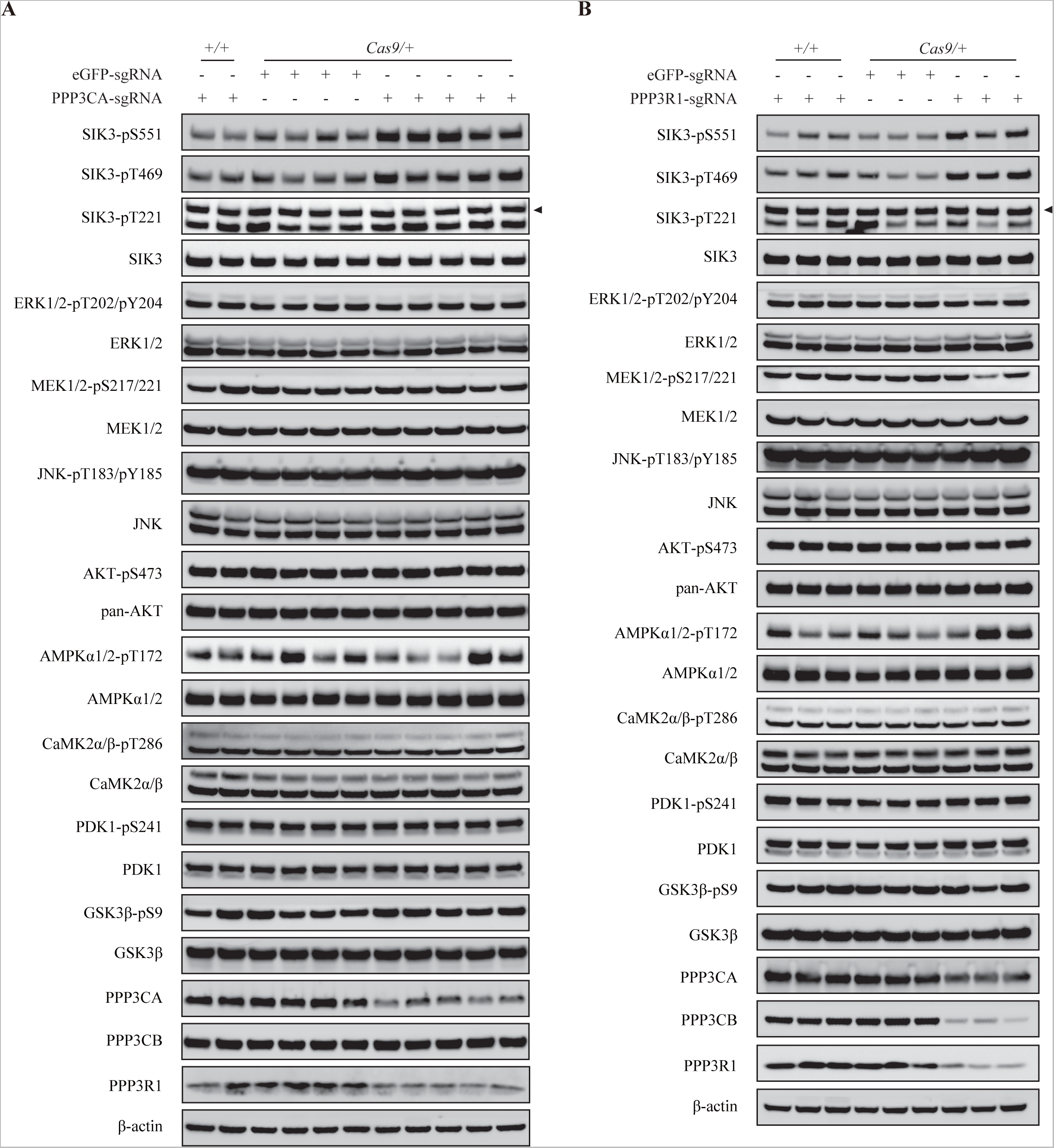
Serine/threonine phosphorylation in the mouse brain after PPP3CA or PPP3R1 knockdown. Phosphorylation of different proteins was examined by anti-phospho-protein antibodies. Each lane shows results from one mouse, and three from each genotype are shown individually here to allow visualization of consistency. **a,** From the left are: two WT mice injected with sgRNAs targeting GFP, five CAS9 expressing mice injected with sgRNAs targeting GFP, and five CAS9 expressing mice injected with sgRNAs targeting PPP3CA. Phosphorylation of SIK3 T469 and SIK3 S551 was reproducibly increased in mice only when sgRNAs targeting PPP3CA were injected into CAS9 expressing mice, neither the WT mice nor with sgRNAs targeting GFP. Phosphorylation of SIK3 T221 did not change. AMPK-T172 phosphorylation was variable in each mouse, not dependent on the expression of CAS9 or sgRNAs targeting PPP3CA. **b,** From the left are: three WT mice injected with sgRNAs targeting GFP, three CAS9 expressing mice injected with sgRNAs targeting GFP, and three CAS9 expressing mice injected with sgRNAs targeting PPP3R1. Phosphorylation of SIK3 T469 and SIK3 S551 was reproducibly increased in mice only when sgRNAs targeting PPP3R1 were injected into CAS9 expressing mice, neither the WT mice nor with sgRNAs targeting GFP. Phosphorylation of SIK3 T221 did not change. AMPK-T172 phosphorylation was variable in each mouse, not dependent on the expression of CAS9 or sgRNAs targeting PPP3R1.

Levels of phosphorylation of different S/T sites of multiple proteins were examined by Western analysis. Phosphorylation levels of T469 and S551 in SIK3 were both increased after PPP3CA or PPP3R1 was knocked down in mouse brains while phosphorylation level of T221 in SIK3 was not significantly changed (Fig. 7 a and b).

We also examined phosphorylation of other protein kinases in the mouse brain. After PPP3CA or PPP3R1 was knocked down, no significant change was consistently observed for the phosphorylation level of specific S/T sites on other proteins such as T202 of ERK1/2, S217 of MEK1, T183 of JNK, S473 of AKT1, T172 of AMPKα, T286 of CaMK2α/β, S241 of PDK1, or S9 of GSK3β (Fig. 7 a and b).

These results provide in vivo evidence that CaN does not dephosphorylate specific S or T in at least 8 other kinases, and that it is specifically required for the dephosphorylation of T469 and S551, but not T221, of SIK3. Our findings that PPP3CA could dephosphorylate S217 of MEK1 precipitated from HEK cells, but that MEK1-S217 phosphorylation was not affected in PPP3CA^KD^ brains, suggest that other PPases are more important than PPP3CA in regulating the phosphorylation of S217 of MEK1 in the brain.

### in vivo Physiologically requirement of PPP3CA for sleep

To investigate physiological roles of the endogenous PPP3CA in mice, we analyzed the sleep phenotypes of PPP3CA^KD^ mice and compared them with those of control (WT^Ctrl^ and eGFP^Ctrl^) mice.

Representative EEG and EMG graphs are shown in Figure 8K and representative hypnogram shown in Figure S8E. There is no significant difference among the three genotypes in terms of the general patterns of EEG or EMG, indicating that PPP3CA^KD^ did not generally or non-specifically disrupt EEG or EMG recordings.

**Fig. 8.**
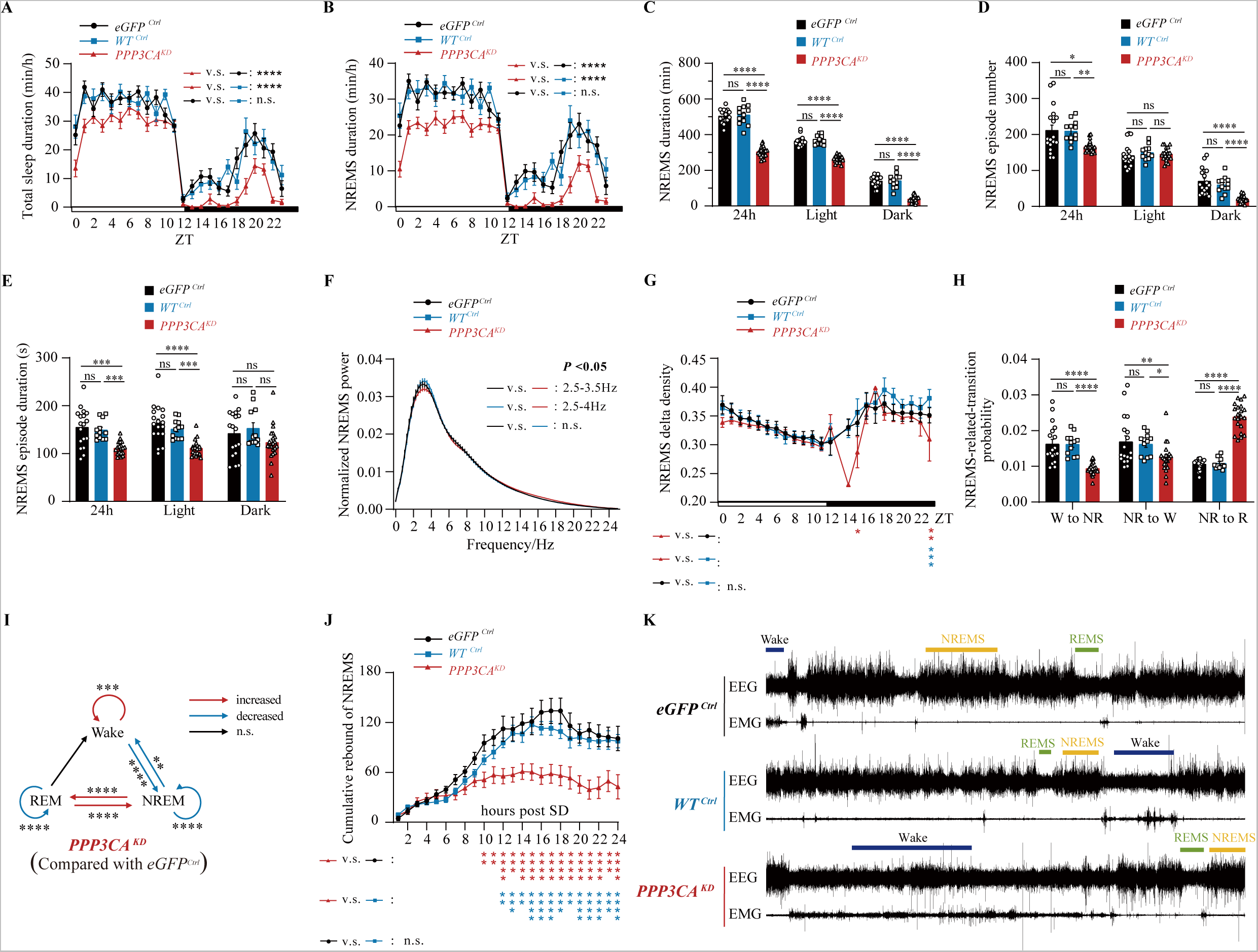
Sleep phenotype of mice after PPP3CA knockdown. Sleep phenotypes of male mice with the genotypes of PPP3CA^KD^ (red, n=21), eGFP^Ctrl^ (black, n=18) and WT^Ctrl^ mice (blue, n=12). PPP3CA^KD^ denotes Rosa26^Cas9/+^ mice injected with AAV-PPP3CA-sgRNA virus; eGFP^Ctrl^ denotes Rosa26^Cas9/+^ mice injected with AAV-eGFP-sgRNA virus; WT^Ctrl^ denotes WT littermate mice injected with AAV-PPP3CA-sgRNA virus. For detailed numbers, see also Extended Data Tables 3 and 4. **a**-**b,** profiles of sleep (**a**) or NREM (**b**). *p< 0.05; ****p<0.0001; ns, not significant; mean±SEM (Two-way ANOVA with Tukey’s multiple comparisons test). **c,** Data and statistics of NREM sleep over 24 hrs. *p< 0.05, ****p<0.0001; ns, not significant; mean ± SEM (One-way ANOVA with Tukey’s multiple comparisons test). **d**-**e**, NREM sleep episode number (**d**) and NREM sleep episode duration of (**e**). *p<0.05; **p<0.01; ***p<0.001; ****p<0.0001; ns, not significant; mean±SEM (Kruskal-Wallis test with Dunn’s multiple comparisons test). **f,** EEG power spectrum during NREM sleep. *p < 0.05; mean ± SEM (Two-way repeated measurement ANOVA with Tukey’s multiple comparisons test). **g,** NREMS delta densities over 14 hrs. X-axis indicates ZT. Data at ZT13 was not included in statistical analysis due to the inaccessible values from PPP3CA^KD^ mice. *p<0.05; **p<0.01; ***p<0.001; ns, not significant; mean±SEM (Mixed-effects model). **h,** Probabilities of transition between different sleep and wake states. *p<0.05; **p<0.01; ***p<0.001; ****p<0.0001; ns, not significant; mean±SEM (Two-way ANOVA with Tukey’s multiple comparisons test). **i,** A diagram illustrating probabilities of transition between different sleep and wke states. **j,** Recovery of NREM sleep after 6 hrs of SD. *p<0.05; **p<0.01; ***p<0.001; ****p<0.0001; ns, not significant; mean±SEM (Two-way repeated measurement ANOVA with Tukey’s multiple comparisons test). **k,** One hr representative EEG and EMG graphs at different vigilance states.

Total sleep over 24 hrs was significantly reduced in PPP3CA^KD^ mice, with reduction at every time point in both daytime (the light phase) and nighttime (the dark phase) (Fig. 8a and Extended Data Fig. 7 a-d, Extended Data Tables 3 and 4). Differences in total sleep over 24 hrs were: 187.6±13.0 mins less in PPP3CA^KD^ than eGFP^Ctrl^ mice, 192.6±14.7 mins less in PPP3CA^KD^ than WT^Ctrl^, 5.0±15.1 mins more in eGFP^Ctrl^ than WT^Ctrl^. In other words, PPP3CA^KD^ mice slept approximately 3 hrs less than either eGFP^Ctrl^ or WT^Ctrl^ control mice. Differences of sleep duration in the 12 hrs of light phase were: 79.5±7.7 mins less in PPP3CA^KD^ than eGFP^Ctrl^ mice, 84.2±8.7 mins less in PPP3CA^KD^ than WT^Ctrl^, 4.8±8.9 mins more in eGFP^Ctrl^ than WT^Ctrl^. Differences of sleep duration in the 12 hrs of dark phase were: 108.1±10.3 mins less in PPP3CA^KD^ than eGFP^Ctrl^ mice, 108.4±11.6 mins less in PPP3CA^KD^ than WT^Ctrl^, 0.2±11.9 mins more in eGFP^Ctrl^ than WT^Ctrl^.

NREM sleep was similar to total sleep in that it was significantly reduced throughout the entire 24 hrs (Fig. 8 b and c, Extended Data Tables 3 and 4). Differences in NREM sleep over 24 hrs were: 201.0±12.4 mins less in PPP3CA^KD^ than eGFP^Ctrl^ mice, 207.0±14.0 mins less in PPP3CA^KD^ than WT^Ctrl^, 5.9±14.4 mins less in eGFP^Ctrl^ than WT^Ctrl^. In other words, NREM was reduced by approximately 204 mins in PPP3CA^KD^ mice as compared to either control. Differences in NREM sleep in the light phase were: 100.8±7.6 mins less in PPP3CA^KD^ than eGFP^Ctrl^ mice, 104.9±8.5 mins less in PPP3CA^KD^ than WT^Ctrl^, 4.1±8.8 mins less in eGFP^Ctrl^ than WT^Ctrl^. During daytime, NREM episode number was not changed (Fig. 8d), but NREM episode duration was reduced (Fig. 8e). Differences in NREM sleep in the dark phase were: 100.2±9.2 mins less in PPP3CA^KD^ than eGFP^Ctrl^ mice, 102.0±10.4 mins less in PPP3CA^KD^ than WT^Ctrl^, 1.8±10.7 mins less in eGFP^Ctrl^ than WT^Ctrl^. During nighttime, the number (Fig. 8d) but not the duration (Fig. 8e) of NREM episodes was reduced.

Changes in REM sleep was much more moderate and were in opposite directions in the light vs the dark phase. Differences in REM sleep over 24 hrs were: 13.3±2.6 mins more in PPP3CA^KD^ than eGFP^Ctrl^ mice, 14.4±2.9 mins more in PPP3CA^KD^ than WT^Ctrl^, 1.0±3.0 mins more in eGFP^Ctrl^ than WT^Ctrl^. In other words, REM was increased by approximately 13 mins in PPP3CA^KD^ mice as compared to either control. Differences in REM sleep in the light phase were: 21.3±2.3 mins more in PPP3CA^KD^ than eGFP^Ctrl^ mice, 20.7±2.6 mins more in PPP3CA^KD^ than WT^Ctrl^, 0.5±2.7 mins less in eGFP^Ctrl^ than WT^Ctrl^ (Extended Data Fig. 7 e and f). In other words, REM sleep was moderately (but statistically significantly) increased during daytime by approximately 20 mins, due to increased REM episode number (Extended Data Fig. 7g) but not REM episode duration (Extended Data Fig. 7h) in PPP3CA^KD^ mice. Differences in REM sleep in the dark phase were: 7.9±1.6 mins less in PPP3CA^KD^ than eGFP^Ctrl^ mice, 6.3±1.8 mins less in PPP3CA^KD^ than WT^Ctrl^, 1.6±1.8 mins more in eGFP^Ctrl^ than WT^Ctrl^ (Extended Data Fig. 7 e and f). In other words, REM sleep was weakly decreased in nighttime by approximately 6 mins (Extended Data Fig. 7 e and f), due to decreased number (Extended Data Fig. 7g) and duration (Extended Data Fig. 7h) of REM episodes.

EEG power spectrum analysis showed only NREM delta power densities reduced in PPP3CA^KD^ mice at ZT15 and ZT23 but not other time points (Fig. 8 e and f). Probabilities of transition between different sleep and wake states were increased between REM and NREM, but decreased between wake and NREM (Figure 8 i and h, Extended Data Fig. 7 k and l).

After 6 hrs of SD, NREM (Fig. 8j) or total sleep/wakefulness (Extended Data Figure 8a) recovered gradually in WT^Ctrl^ and eGFP^Ctrl^ mice, whereas their recoveries after SD was significantly reduced in PPP3CA^KD^ mice. Recovery of REM after SD was not significantly different among WT^Ctrl^, eGFP^Ctrl^ and PPP3CA^KD^ mice (Extended Data Fig. 8b).

Thus, PPP3CA is physiologically required for basal sleep mainly by promoting NREM episode duration during daytime and NREM episode number during nighttime. PPP3CA regulation of REM was moderate and different between daytime and nighttime. PPP3CA is also physiologically required for recovery of NREM after SD.

### in vivo Physiologically requirement of PPP3R1 for sleep

PPP3CA is a catalytic subunit of the CaN phosphatase. Its involvement in sleep naturally begs the question whether a regulatory subunit is also involved. Because PPP3R2 was specifically expressed in the testis, not in the brain, we investigated PPP3R1 for potential roles in regulating sleep.

Western analysis of whole brains showed that PPP3R1^KD^ mice expressed PPP3R1 protein at a significant decreased level (Fig. 7b). Representative EEG and EMG graphs are shown in Fig. 9k and representative hypnograms in Extended Data Fig. 10e.

**Fig. 9.**
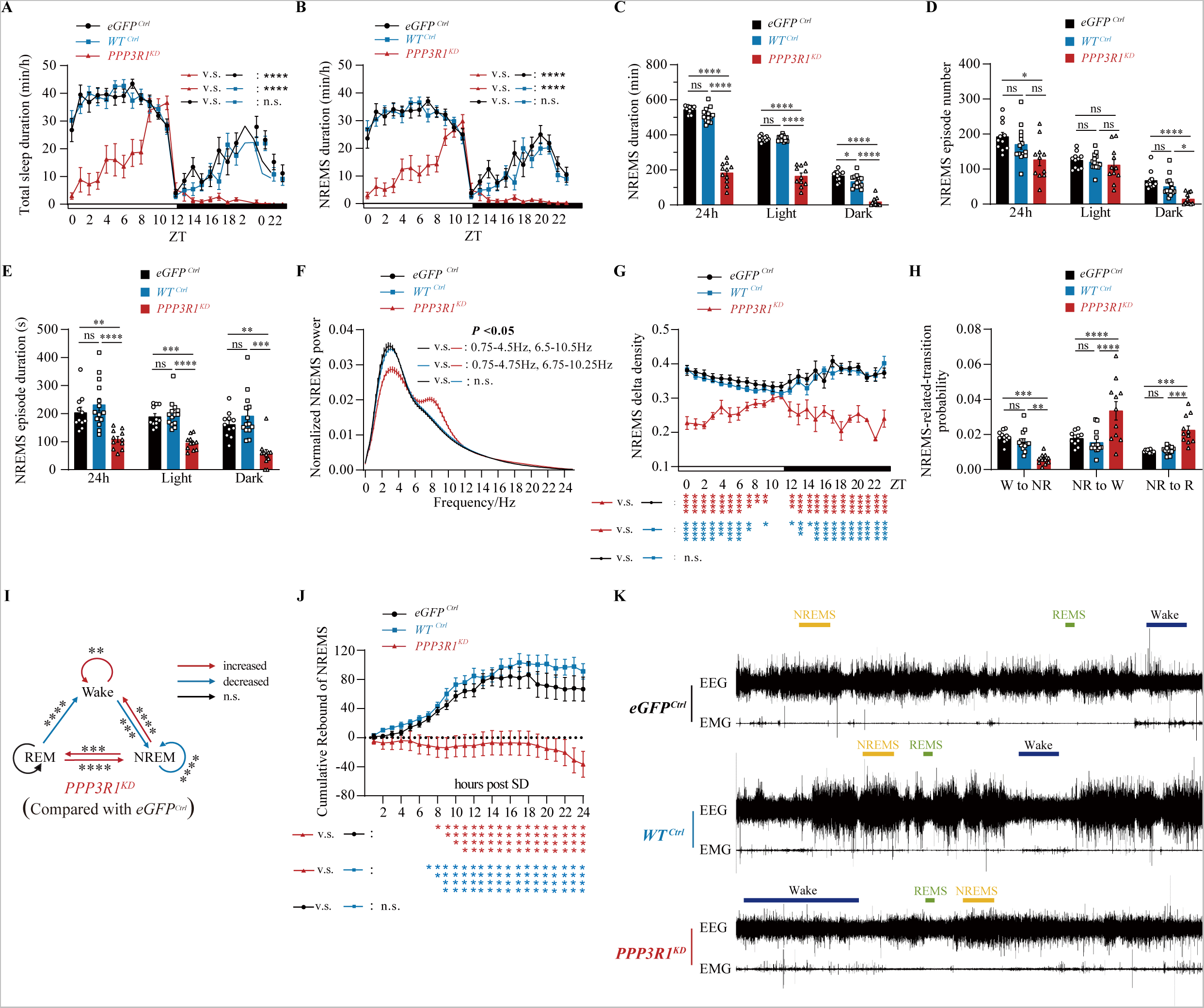
Sleep phenotype of mice after PPP3R1 knockdown. Sleep phenotypes of male mice with the genotypes of PPP3R1^KD^ (red, n=11), eGFP^Ctrl^ (black, n=11) and WT^Ctrl^ mice (blue, n=14). eGFP^Ctrl^: Rosa26^Cas9/+^ mice injected with AAV-eGFP-sgRNA virus. WT^Ctrl^: WT mice injected with AAV-PPP3R1-sgRNA virus. PPP3R1^KD^: Rosa26^Cas9/+^ mice injected with AAV-PPP3R1-sgRNA virus. For details, see also Extended Data Tables 5 and 6. **a**-**b,** Sleep profiles (**a**) or profiles of NREM sleep (**b**). ns: statistically not significant; ****p<0.0001; mean ± SEM (Two-way ANOVA with Tukey’s multiple comparisons test). **c,** Data and statistics of NREM sleep over 24 hrs. ns: statistically not significant; *p< 0.05; ****p<0.0001; mean ± SEM (One-way ANOVA with Tukey’s multiple comparisons test). **d**-**e,** NREM sleep episode number (**d**), and NREM sleep episode duration (**e**). ns: statistically not significant; *p< 0.05; **p<0.01; ***p<0.001; ****p<0.0001; mean ± SEM (Kruskal-Wallis test with Dunn’s multiple comparisons test). **f,** NREMS EEG power spectrum analysis. X-axis indicates frequency distribution of EEG power. *p<0.05; mean ± SEM (Two-way repeated measurement ANOVA with Tukey’s multiple comparisons test). **g,** NREMS delta power density over 24 hours. X-axis indicates ZT. Statistical analysis each hr shown in *s. ns: statistically not significant; *p< 0.05; **p<0.01; ***p<0.001; ****p<0.0001; mean ± SEM (Mixed-effects model). **h,** Transition probabilities of different sleep and wake states. W: wake, NR: NREM Sleep, R: REM Sleep. ns: statistically not significant; *p< 0.05; **p<0.01; ***p<0.001; ****p<0.0001; mean ± SEM (Two-way ANOVA with Tukey’s multiple comparisons test). **i,** A diagrammatic illustration of transition probabilities of different sleep and wake states, summarized from data in Fig. 9H and Fig. S9K and S9L. ns: no significant changes. **j,** Cumulative NREMS rebound after 6 hrs of SD. ns: statistically not significant; *p< 0.05; **p<0.01; ***p<0.001; ****p<0.0001; mean ± SEM (Two-way repeated measurement ANOVA with Tukey’s multiple comparisons test). **k,** Representative one hour EEG and EMG signals.

Sleep duration at every ZT in both light and dark phases except 5 ZT hrs around light to dark transition was significantly and dramatically decreased in PPP3R1^KD^ mice (Fig. 9a and Extended Data Fig. 9 a-d, Extended Data Tables 5 and 6). PPP3R1^KD^ mice spent significantly reduced amount of time for sleep over 24 hrs: 236.4±24.5 mins for PPP3R1^KD^ mice (Extended Data Table 5). This is 381.4±22.8 mins less than eGFP^Ctrl^ mice or 349.3±21.5 mins less than WT^Ctrl^ mice (Extended Data Table 6). In other words, sleep over 24 hrs is decreased by more than 5 hrs in PPP3R1^KD^ mice.

Over 24 hrs, NREM was reduced from 543.6±6.5 mins in eGFP^Ctrl^ or 512.0± 0.6 mins in WT^Ctrl^ to 183.6±18.3 mins in PPP3R1^KD^ mice (Fig. 9c, Extended Data Table 5). This is 360.1±18.5 mins less in PPP3R1^KD^ than eGFP^Ctrl^ mice or 328.4±17.5 mins less than WT^Ctrl^ mice (Extended Data Table 6). The decrease in total NREM over 24 hrs was attributable to shorter NREMS episode duration (Fig. 9c) but not to episode number (Fig. 9d).

In the light phase, NREM sleep (Figs 9 b-e; Extended Data Tables 5 and 6) was significantly decreased in PPP3R1^KD^ mice whereas REM was not significantly different between PPP3R1^KD^ and eGFP^Ctrl^ mice or PPP3R1^KD^ and WT^Ctrl^ mice (Extended Data Fig. 9 e-h; Extended Data Tables 5 and 6). NREM of PPP3R1^KD^ mice in the light phase is 212.9±14.5 mins less than eGFP^Ctrl^ mice or 213.4±13.7 mins less than WT^Ctrl^ mice (Extended Data Table 6). In the dark phase, sleep of any kind (including both REM and NREM) was dramatically reduced in PPP3R1^KD^ mice: 20.4±8.5 mins total sleep, 18.47±7.7 mins of NREM sleep, 1.9±0.9 mins of REM sleep (Extended Data Table 5), corresponding to a reduction of 159.0±14.1 mins less than eGFP^Ctrl^ mice or 123.8±13.3 mins less than WT^Ctrl^ mice, 147.2±13.2 mins less than eGFP^Ctrl^ mice or 115.0±12.5 mins less than WT^Ctrl^ mice, 11.7±2.0 mins less than eGFP^Ctrl^ mice or 9.0±1.8 mins less than WT^Ctrl^ mice (Extended Data Table 6). These results indicate that PPP3R1 is physiologically required for sleep.

Probabilities of transition between different sleep and wake states (Fig. 9i) were summarized from data in Figure 9H and Figure S9K and S9L. Probabilities of transition from Wake to NREMS, and from NREMS to NREMS, were reduced whereas probabilities of transition from NREMS to wake, wake to wake were increased.

PPP3R1^KD^ mice showed neither NREMS nor REMS sleep recovery after 6 hrs of SD, with statistically significant decreases at every time point tested from the 8^th^ to the 24^th^ hr (Fig. 9j and Extended Data Fig. 10 a-b).

NREM delta power densities were decreased in PPP3R1^KD^ mice (Fig. 9 f-g and Extended Data Fig. 9 i-j), suggesting that PPP3R1 is required for sleep need. After SD, NREMS delta power recovery was also less in PPP3R1^KD^ mice than control mice (Extended Data Fig. 10 c-d).

## Discussion

We conclude that CaN plays a major role in controlling sleep, especially of the NREM type, and suggest a signaling pathway involving not only protein kinases but also protein phosphatases in sleep regulation. Our results have also demonstrated the specificity of CaN in phosphorylating specific S and T sites in SIK3, revealing a specificity previously unsuspected for PPases.

Of the three catalytic subunits of CaN, all are active in vitro in biochemically dephosphorylating T469 and S551, but not T221, of SIK3. in vivo, we have proven a biochemical role of dephosphorylating T469 and S551 for PPP3CA and PPP3CB in HEK cells and for PPP3CA in the mouse brain. We have found a physiological role for PPP3CA in regulating sleep in the mouse. PPP3CB is also involved in HEK cells in mediating dephosphorylation induced by Ca^2+^, but it remains to be tested whether PPP3CB is required in the brain, where it is also expressed, though at a level lower than that of PPP3CA. The finding that the effect of PPP3R1 knockdown in the mouse was stronger than PPP3CA knockdown, especially on NREM sleep, supports the possibility that PPP3R1 also functions with a catalytic subunit other than PPP3CA. PPP3CB provides the obvious candidate. The expression of PPP3CC is thought to be testis-enriched or -specific, and its role in sleep has not been tested. Of the two regulatory subunits, we have shown biochemically that PPP3R1 is required for SIK3 dephosphorylation in HEK cells and in the mouse brain, and physiologically that PPP3R1 is required for sleep regulation. We have not targeted PPP3R2 in the mouse because it is testis-specific. Because of the interdependence of catalytic and regulatory subunits (Fig. 7 a-b), the sleep phenotypes observed by us in the manuscript should be attributed to CaN, not distinguishing among different subunits.

Is SIK3 the only downstream target mediating the physiological role of CaN in sleep regulation? It is not clear at the moment. The possibility that CaN may involve substrates in addition to SIK3 is supported by the following: the sleep phenotypes of PPP3CA^KD^ and PPP3R1^KD^ were stronger than that of SIK3^T469A/T469A^ mutant mice, which could be explained by either more sites in SIK3 for PPP3CA dephosphorylation, or by more substrates in addition to SIK3 for PPP3CA dephosphorylation. SIK 1, 2, 3 are all members of the AMPK related kinases (ARKs), we are actively examining whether and which one of the 16 to 20 ARKs ^87–89^ is involved in sleep. Direct effects of CaN on targets other than ARKs are also being explored.

### Advantage of combining classic biochemistry with new chemical biology

We have used both a classic biochemical purification method and a new chemical biological photo-crosslinking method. This combination is very helpful. With the biochemical method (Figs. 2 and 3), 4 PPases were purified. It was unclear which one among those 4 was the right phosphatase and it was also possible that there were other PPases. With the photo-crosslinking method, PPases were not the most prominent proteins found to interact with SIK3-HA in the brain. However, when we looked at both results, it immediately drew our attention to one PPase: the one shared by both methods.

The combination thus expediated the process of focusing on CaN as the PPase for T469 and S551 of SIK3.

### A new photo-crosslinking strategy

We (RG and PRC) have developed a new photo-catalytic crosslinking strategy for capturing in situ protein-protein interactions in native tissue slice samples (Fig. 4). Our method utilizes eosin as photocatalyst which generates singlet oxygen upon visible light irradiation. The singlet oxygen activates a diamine probe to capture direct protein-protein interactions. Proteins of interest (POIs) and their interacting partners could then be pulled down and analyzed by mass spectrometry.

This method has several unique advantages: 1) it does not require transfection or other genetic methods to introduce crosslinking probes. Ready-to-use in native samples and captures in situ protein-protein interactions; 2) the photocatalyst eosin generates singlet oxygen with high efficiency under irradiation. The molecule itself also performs high compatibility in tissue samples as a tissue staining dye; 3) the peak excitation wavelength of eosin is around 520 nm. Visible light-meditated crosslinking method shows benefit against traditional UV methods with better tissue penetration and less cytotoxicity.

### Specificity of CaN dephosphorylation of sites in SIK3

With the background of the widely held perception of PPases being not very specific, the selectivity of a PPase for different sites in the same substrate protein was striking.

We have three pieces of in vitro evidence supporting the specificity of CaN (PPP3CA and PPP3R1): 1) PPP3CA expressed in and purified by immunoprecipitation from HEK cells dephosphorylated T469 and S551, but not T221, of SIK3 (Fig. 5); 2) With each protein component expressed in and purified from *E. coli*, we have demonstrated that dephosphorylation of T469 and S551 in SIK3 requires Ca^2+^, CaM, PPP3CA and PPP3R1 (Fig. 5f); 3) The same purified components combined together did not dephosphorylate T221 of SIK3. CaN also did not dephosphorylate T308 of AKT1, S241 of PDK1, T215 of MARK1, S9 of GSK3β, T183 of JNK1, or T185 of ERK2, though it dephosphorylated S217 of MEK1 (Fig. 5 b-e).

We have three pieces of in vivo evidence supporting the specificity of CaN: 1) Overexpression of CaN in HEK cells enhanced Ca^2+^ ionophore induced dephosphorylation of T469 and S551 in SIK3, but did not affect phosphorylation of T221 in SIK3 (Fig. 6 c-d); 2) Knockdown of the catalytic or the regulatory subunits of CaN in HEK cells inhibited the T469 and S551 dephosphorylation induced by Ca^2+^ ionophore (Fig. 6e). Two subunits of CaN (PPP3CA and PPP3CB) had to be targeted by sgRNAs at the same time to suppress the dephosphorylation effective and robustly, suggesting that, in HEK cells, both catalytic subunits function in mediating the ionophore response. 3) This is not the case in the mouse brain where PPP3CA is highly expressed. Knockdown of PPP3CA alone was sufficient to inhibit the dephosphorylation of T469 and S551 of SIK3 (Fig. 7a). PPP3CA knockdown did not affect the phosphorylation level of T221 of SIK3, proving the site-specificity of CaN for SIK3. PPP3CA or PPP3R1 knockdown in the mouse brain also did not affect the phosphorylation of specific sites in kinases such as ERK1/2, MEK1, JNK, AKT, CaMK2α/β, PDK1, or GSK3β (Fig. 7 a-b).

The specificity of CaN for T469 and S551 of SIK3 raises the question whether there is a site specific PPase for T221 of SIK3. More broadly, are there kinase-phosphatase pairs in all important signaling pathways?

### CaN regulation of sleep

PPP3CA knockdown reveals that it is required physiologically for regulating sleep both during daytime and nighttime, especially NREM (Fig. 8 a-e). Its role in REM sleep is moderate and different between daytime and nighttime (Extended Data Fig. 7 e-h). Different from LKB1^17^ or SIK3^19^ knockout mice, sleep need measured by NREM delta power densities was not affected by PPP3CA knockdown (Fig. 8 f and g). PPP3CA is required for recovery of NREM sleep (Fig. 8j), but not REM sleep (Extended Data Fig. 8b), after deprivation, indicating that PPP3CA is important for homeostatic regulation of sleep.

PPP3R1 knockdown showed a stronger phenotype than PPP3CA, which is expected because a regulatory subunit can interact with all catalytic subunits in the brain, while, although the catalytic subunit could also interact with all regulatory subunit, there is only one regulatory subunit in the mouse brain, with the other one (PPP3R2) limited to the testis.

PPP3R1 is required physiologically for both NREM and REM sleep. It is required for nighttime sleep, and its reduction led to near elimination of NREM and REM sleep at nighttime (Fig. 9 a-c; Extended Data Fig. 9 e-f). During daytime, it is required more for NREM sleep (Fig. 9 b-c; Extended Data Fig. 9 e-f). It is required for sleep need indicated by NREM delta power densities (Fig. 9g), and for recovery of both NREM (Fig. 9j) and REM (Extended Data Fig. 10b) after SD.

### Calcium regulation of sleep

Ca^2+^ imaging in different brain regions have shown that intracellular and extracellular concentrations of Ca^2+^ are different among different sleep/wake states^90–93^. Pharmacological and genetic manipulations of channels affecting Ca^2+^ concentrations could change sleep patterns in mice^38, 94–97^.

Our findings of the roles of CaN provides one possible downstream component of Ca^2+^, but it remains to be further studies how Ca^2+^, kinases, phosphatases, ion channels and transcriptional factors interact in sleep regulation.

### Protein kinases and sleep

The first kinase implicated in sleep regulation was PKA. An antidepressant capable of inhibiting phosphodiesterase and increasing cyclic adenosine monophosphate (cAMP) could increase wakefulness^26^ in rats. In mice, overexpression of a dominant negative form of PKA increased REM sleep and NREM fragmentation while decreasing sleep rebound after deprivation^30^. ERK requirement for sleep in mice was shown by significant sleep reduction after pharmacological inhibitions or genetic deletions of ERK1 or ERK2 genes in neurons^33^. AMPK was implicated in sleep regulation when its inhibitor was found to decrease sleep and its activator found to increase sleep in mice^34^. Pharmacological inhibition of CaMKII in specific brain regions was found to increase sleep^37^. Genetic knockout studies show the importance of CaMK2α and CaMK2β in sleep with a reduction of approximately 50 minutes (mins) or 120 mins per 24 hrs in CaMK2α and CaMK2β knockout mutants, respectively^38^. SIK3 was discovered through a forward genetic screen in mice when a gain of function mutant of SIK3 was discovered^14^. A small fragment was deleted in the SIK3^sleepy^ mutant, resulting in the absence of a PKA target site S551^20^. Mutations of the S551 equivalent sites in SIK1 and SIK2 led to the GOF phenotype^21^, making it unclear which of the SIK kinases or phosphorylation sites were physiologically required for sleep regulation. Our studies of mice deleting each one of the SIK genes indicated that only SIK3, but not SIK1 or SIK2 is required for sleep in mice^19^. LKB1 is a tumor suppressor gene^41–43, 98, 99^ whose product was found to phosphorylate T172 of the α subunit of AMPK^100–106^ and the equivalent site in ARKs including SIK3^87^. Our recent functional studies in vivo have shown requirement of LKB1 for sleep in both flies and mice^17^. The simple scenario of LKB1 upstream of SIK3 in sleep regulation was rendered uncertain by our in vitro biochemical studies which has uncovered more than twenty kinases of the STE20 subfamily in addition to LKB1 upstream of ARKs^107,108^. We are still studying whether any of the STE20 kinases identified by us biochemically is involved in sleep regulation.

Our discovery of specific phenotypes for CaN in sleep will stimulate further studies of other PPases in regulating sleep. For example, while it seems that CaN is more important for NREM than REM, would there be a PPase more important for regulating REM sleep? What are the relationships between kinases and phosphatases in sleep regulation? The specificities we discovered for CaN in dephosphorylating specific sites and in regulating specific components of sleep brings both questions and excitement for furthering our understanding of molecular mechanisms of mammalian sleep. More broadly, specific involvement of PPases in important physiological processes should be further studied.

## Methods

### Antibodies

The following primary antibodies were used: anti-HA tag (C29F4, CST), anti-FLAG M2 HRP conjugated (A8592, Sigma), anti-SIK3 (Santa Cruz, sc-515408), anti-SIK3 pT221 (Abcam, ab271963), anti–SIK3 pT469 (Abcam, ab225633), anti–SIK3 pS551 (Abcam, ab225634), anti-PPP3CA (ABclonal, A1063), anti-PPP3CB (AffinitY, DF12705), anti-PPP3CC (ABclonal, A7714), anti-PPP3R1 (ABclonal, A0954), anti-JNK (CST, 9252), anti-phospho-JNK (CST, 9251), anti-ERK1/2(CST, 4695), anti-phospho-ERK (CST, 4370), anti-AMPKα1/α2 (ABclonal, A12718), anti-AMPKα1/2– pT183/T172 (ABclonal, AP1345), anti-PDK1 (ABclonal, A8930), anti-PDK1 pS241 (ABclonal, AP1357), anti-MEK1/2 (ABclonal, A4868), anti-MEK1/MEK2 pS217/S221 (ABclonal, AP1349), anti-GSK3β (CST, 12456), anti–GSK3β pS9 (CST, 5558), anti-panAkt (CST, 4685), anti-Akt pS473 (CST, 4060), anti-CaMKII pT286 (Abcam, ab171095), anti-CaMKIIα/β (CST, 4436S), anti–β-actin (Abcam, ab8226).

### Cell culture and cDNA transfection

HEK293T cells were cultured in Dulbecco’s modified Eagle’s medium (DMEM, Gibco) medium containing 10% fetal bovine serum (FBS, Gibco) and 1% Penicillin/Streptomycin (Gibco). cDNAs were transfected into HEK293T cells with Lipofectamine 3000 reagent (Thermo Fisher) according to the manufacturer’s instructions and harvested 24 to 28 hrs after transfection.

### Drug treatment and protein preparation

HEK293T cells were treated with ionomycin (MedChemExpress) at indicated concentrations and time durations at 37 ℃. Cells were then harvested and lysed with 1 ml lysis buffer (0.3% Chaps, 10 mM KCl, 1.5 mM MgCl_2_, 1 mM EDTA, 1 mM EGTA, pH 7.4, 1x protease inhibitor cocktail (Roche), 1x phosphatase inhibitor II and 1x phosphatase inhibitor III (Sigma)) before centrifugation of cell lysates at 13000 rpm for 10 min at 4 ℃. Protein concentrations of cell lysates were determined with the bicinchoninic acid (Thermo Fisher, 23225) assay and normalized to 2 mg/ml. Samples were analyzed by immunoblotting with the indicated antibodies.

### Mice

WT C57 BL/6J mice (8 to 10 weeks old) were purchased from Beijing Vital River Laboratories Technology Co., Ltd. or Laboratory animal resource center in Chinese Institute for Brain Research. Rosa26-Cas9 knock-in (Rosa26^Cas9/+^, RRID: IMSR_JAX:024858) mice were obtained from Jackson laboratory ^109^. SIK3-3xHA mice were constructed in BIOCYTOGEN by infusing a 3xHA-T2A-iCre cassette into the C terminus right before the SIK3 stop codon. All experimental procedures were performed in accordance with the guidelines and were approved by the Animal Care and Use Committee of Chinese Institute for Brain Research, Beijing. Mutant mice and wt littermates were maintained on a C57 BL/6J background. Mice were housed under a 12 hr:12 hr light/dark cycle and controlled temperature and humidity conditions. Food and water were delivered *ad libitum*. Mice used in all experiments were 10-14 weeks old.

### Generation of mice with point mutations in SIK3

SIK3 point mutant mice were constructed with CRISPR-Cas9 mediated homologous recombination. For SIK3^T469A/T469A^, the gRNA sequence was 5’-TTTGTCAATGAGGAGGCACA-3’ and a single strand homologous arm was designed to introduce nucleotide mutation from ACG to GCG (T469A) as well as a restriction enzyme site *BstUI* for future genotyping, sequence of which was 5’-CCTTCTCCAGAAGCCTTGGTTCGCTATTTGTCAATGAGGAGGCACGCGGTGGGAGTGGCTGACCCACGGTAAGTACCTGGTCAGCATCCT-3’. A mixture of Cas9-expressing mRNA, single strand homologous arm and sgRNA was injected into fertilized eggs through electroporation and the eggs were then transplanted into the womb of foster mothers. F0 and F1 mice were genotyped through PCR and *BstUI* digestion to make sure the presence of recombination. Mutant lines were back-crossed to C57BL/6J for at least 5 generations to exclude possible off-targeting.

### Mouse brain protein preparation

Whole brains of mice were quickly dissected, rinsed withP BS and homogenized by homogenizer (Wiggens, D-500 Pro) in ice-cold lysis buffer (150 mM NaCl, 1% Triton-X-100, 0.5% sodium deoxycholate, 0.1% SDS, 50 mM Tris-base, freshly supplemented with a protease and phosphatase inhibitor cocktail). Brain homogenates were centrifuged at 15000 rpm for 25 min at 4 ℃. Supernatants were carefully transferred into a new microtube. Protein concentrations of brain lysates were determined with the bicinchoninic acid assay and normalized to 2 mg/ml. Before immunoblotting, samples were kept in liquid nitrogen, if necessary.

### Biochemical purification

Lysates from HEK 293 cells were prepared and filtered through 0.45 μm filters. 500 mg cell lysates at a concentration of 10 mg/ml were fractionated on a QHP anionic chromatography column, eluted with a linear gradient of NaCl (0-600 mM) into 20 column volumes (CVs), with each fraction collected as one CV as samples 1 to 20, and the final wash with 1 M NaCl buffer A with 5 CVs gave rise to samples 21 to 25. Each fraction was dialyzed into buffer A, removing NaCl. 10 μl of each sample was used for analysis of activities removing phosphate from SIK3 T469 and S551. T469 and S551 of bacterially expressed recombinant SIK3 were phosphorylated by PKA in vitro before being used to test phosphatase activities of the fractions of HEK lysates. Fractions 10 to 13 contained significant activities removing phosphate from SIK3 T469 and S551.

Fractions 10 to 13 from QHP were combined and dialyzed with buffer A before being loaded onto a Blue HP column. It was eluted with a linear gradient of NaCl (0-600 mM) into 20 column volumes (CVs), with each fraction collected as one CV as samples 1 to 20, and the final wash with 1 M NaCl buffer A with 5 CVs gave rise to samples 21 to 25. The FL fraction from the Blue HP contained significant activities removing phosphate from SIK3 T469 and S551.

The FL fraction from the Blue HP column was dialyzed with buffer A and loaded onto a SPHP column. The FL fraction from the SP-HP column contained significant activities removing phosphate from SIK3 T469 and S551.

The FL fraction from the SP-HP column was dialyzed with buffer A and loaded onto a heparin HP column. The FL fraction and Fraction 1 contained significant activities removing phosphate from SIK3 T469 and S551.

The FL fraction from the heparin HP column was dialyzed with buffer A and loaded onto a HAPHP column. The rest of the fractionation was similar to the first column except that the final wash was with 5 CVs of 500 mM K_2_PO_4_, giving rise to samples 21 to 25. Fractions 5 to 7 contained significant activities removing phosphate from SIK3 T469 and S551.

Active fractions from the HAPHP column were condensed into 0.5 ml, fractionated on a Superdex 200 molecular sieve column, eluted with 200 mM NaCl gradient into 20 CVs. 1 ml from each fraction was collected and labeled as samples 1 to 20. Protein contents were monitored with UV at 280 nm.

### Incorporation of photocatalytic probes into mouse brain slices for visible light-induced PPI capture

SIK3-3xHA mice were sacrificed and brain slices were kept in PBS. Neurobasla medium (Gibco) supplied with B-27 for photocatalytic crosslinking was prepared in darkness by adding eosin Y and 1,6-dihexamine to a working concentration of 50 μM and 1 mM, respectively. Cell culture chambers (Millipore, PICM0RR50) were placed in 6-well plates and rinsed with PBS. Brain slices were transferred to chambers carefully with a sterile dropper and each chamber finally contained four slices to ensure thorough stretch of each slice. PBS was discarded by pipetting from the outer side of chambers and 0.5 ml of neurobasal medium containing photocatalytic crosslinkers was added into each well to ensure that each slice was totally infiltrated with the probes. Samples were incubated at 37°C with 5% CO_2_ for 1 hr before photo-irradiation. For photocatalytic crosslinking, the plates were placed on green LED (520 nm, 20 mW/cm^2^) equipment while an ice bag and a fan were used to reduce light irradiation-induced heat. After 15 mins of green light irradiation, the color of the medium was bleached, indicating effective activation of the eosin probe. The medium was discarded and after one round of PBS wash, the slices was collected into 1.5 ml Eppendorf tubes, which were placed into liquid N_2_ to freeze the slices.

### Enrichment of crosslinked proteins

To each tube with frozen brain slice samples was added 1 ml of ice-cold lysis buffer containing 1% of protease cocktail and a steel ball. Samples were lysed through ultrasonication. After centrifugation (12,000 g, 10 mins, 4 °C) to discard the residue, crosslinked proteins were enriched by anti-HA magnetic beads (Pierce, 88837) according to the manufacture protocol. Importantly, the beads should be washed with lysis buffer (with 0, 0.25, and 0.5 M NaCl) for three times to diminish non-specific binding proteins. Crosslinked proteins were eluted with 2x SDS-loading buffer and heated to 95 °C for 10 min. Eluted proteins were subjected to further Western analysis and LC-MS/MS.

### Protein digestion and dimethyl labeling

#### In-gel digestion

Proteins enriched with HA beads were loaded on an 8% SDS-PAGE gel and run at 150 V for 30 mins. After silver staining, the desired bands of protein mixture were excised and cut into 1 mm^3^ pieces. The gel pieces were discolored in discoloring buffer until they all turned transparent. A dehydration process was carried out by adding pure acetonitrile into gel pieces until they were totally dehydrated to appear non-transparently white. Samples were then incubated in the reduction buffer (10 mM DTT, 50 mM ammonium bicarbonate) at 56 °C for 30 mins and further incubated in alkylation buffer (55 mM iodoacetamide, 50 mM ammonium bicarbonate) at 37 °C for half an hr in the dark. After washed by 50 mM ammonium bicarbonate buffer twice, gel pieces were dehydrated through the same protocol. 20 ng/μl trypsin buffer in 50 mM ammonium bicarbonate was added and samples were incubated at 4 °C for one hr. The remaining buffer was discarded and 50 mM ammonium bicarbonate buffer was added for another 16 hrs of digestion at 37 °C. The resulting peptides were extracted with extraction buffer (50% acetonitrile, 45% water and 5% formic acid), before being centrifuged to dryness under vacuum.

#### Dimethyl labeling

The collected peptides were reconstituted in 100 mM TEAB buffer. 39.688 mg/mL of NaBH_3_CN followed by 4% (v/v) CH2O or CD2O were added for light and heavy dimethyl labeling, respectively, following the addition of 39.688 mg/ml NaBH_3_CN. After incubation in a fume hood for 30 mins at room temperature, enough 1% (v/v) ammonia solution and FA were added immediately to quench the labeling reaction. The light and heavy samples were combined together, and then desalted and dried under vacuum.

#### LC-MS/MS analysis

Trypsin digested peptides were analyzed on a Exploris 480 Hybrid Quadrupole Orbitrap Mass Spectrometer as well as Thermo Scientific Q Exactive Orbitrap Mass Spectrometer in conjunction with an Easy-nLC II HPLC (Thermo Fisher Scientific). The mobile phases were A: 0.1% formic acid in H_2_O; B: 0.1% formic acid in 80% ACN–20% H_2_O. MS/MS analysis was performed under the cationic mode with a full-scan *m*/*z* range from 350 to 1,800 and a mass resolution of 70,000.

#### Peptides identification

For quantitative SIK3 interactome analysis, the quantification of light/heavy ratios was calculated with a precursor mass tolerance of 20 ppm. Alkylation of cysteine (+57.0215 Da) was set as the static modification, and oxidation of methionine (+15.9949 Da) and acetylation of N-terminal Lys (+42.0106 Da) was assigned as the variable modification. The isotopic modifications (28.0313 and 32.0557 Da for light and heavy labeling, respectively) were set as fixed modifications on the peptide N-terminus and lysine residues. Half-tryptic terminus and up to two missing cleavages were set within tolerance.

#### Co-immunoprecipitation

For HEK293T cells, plasmids expressing FLAG-tagged SIK3 and HA-tagged PPP3CA were transfected for 24 hrs. Cells were then collected and lysed with 1 ml lysis buffer (25 mM Tris-base pH 7.4, 150 mM NaCl, 1% NP40, 1mM EDTA, 5% glycerol,1x protease inhibitor cocktail, 1x phosphatase inhibitor II and 1x phosphatase inhibitor III) before centrifugation at 13000 rpm for 10 min at 4℃. 40 μl supernatants were transferred into new microtubes as input samples and the rest was incubated either with 20μl anti-FLAG (Millipore, M8823) or anti-HA antibody coated magnetic beads balanced by lysis buffer for 1 hr at 4 ℃. Beads were then washed with 1 ml lysis buffer three times at 4 ℃ and 40 μl PBS was added to transfer the beads into new microtubes as enriched samples. For mouse, anti-PPP3CA or anti-SIK3 antibody and corresponding IgG were pre-incubated with protein A/G beads (YEASEN) for 2 hrs at 4℃ in lysis buffer. Protein samples from mouse brain were at first pre-absorbed using IgG-protein A/G beads for 30 min at RT and 40 μl supernatants were transferred into a new microtube as input sample before anti-PPP3CA/SIK3 protein A/G beads was added and rotated overnight at 4℃. Beads were then washed with 1 ml lysis buffer three times at 4℃ and 40 μl PBS was added to transfer the beads into a new tube as enriched samples.

### Expression of recombinant proteins in *E. coli*

cDNAs for specific proteins were subcloned into the pET-28a vector, with appropriate tags such as MBP, GFP or FLAG. Plasmids were transfected into *E. coli* BL21 and incubated at 37 ℃ until the OD reached 0.6, when 0.5 mM of IPTG was added at 18 ℃ to induce protein expression for 16 hrs. Cells were collected and treated with Ni column binding buffer (300 mM NaCl, 20 mM Tris-HCl, pH 7.5) containing protease inhibitors and thoroughly suspended. Cells were lysed with ultrasonication before centrifugation (14,000 rpm, 30 mins, 4℃). Supernatants were filtered through 0.45 μm and purified by Ni beads to 90% purity. Eluted proteins were measured with Coomasie blue and the rest of the proteins were stored at −80 ℃.

### in vitro Phosphatase assay

Plasmids expressing FLAG-tagged SIK3 were transfected into HEK293T cells for immunoprecipitation. After 24 hrs, cells were collected and lysed with 1 ml lysis buffer (25 mM Tris-base pH 7.4, 150 mM NaCl, 1% NP40, 1mM EDTA, 5% glycerol,1x protease inhibitor cocktail, 1x phosphatase inhibitor II and 1x phosphatase inhibitor III) before centrifugation at 13000 rpm for 10 min at 4 ℃. Cell lysates were incubated with 20μl anti-FLAG antibody coated magnetic beads balanced by lysis buffer for 1 hr at 4 ℃. Beads were washed with 1 ml lysis buffer three times at 4 ℃, before final elution with 30 μl buffer A containing 2 mg/ml 3xFLAG peptide. Phosphatase reactions were performed for 2 hrs at 37℃ by adding 4 μl FLAG-SIK3, 3 μg rPPP3CA, 3 μg rPPP3R1, 4 μg rCaM with a final concentration of 10 mM CaCl_2_ and 20 mM MgCl_2_. Samples were analyzed by immunoblotting with the indicated antibodies.

### Viruses

Viruses used in this study: AAV2/PHP.eB-CMV-mScarlet-PPP3CA-sgRNA-WPRE, AAV2/PHP.eB-CMV-mScarlet-PPP3R1-sgRNA-WPRE and AAV2/PHP.eB-CMV-mScarlet-eGFP-sgRNA-WPRE.

PPP3CA^KD^ and PPP3R1^KD^ mice were generated with triple-targeted CRISPR-Cas9 technology ^110^ by virus injection. Plasmids were generated before virus package. Three sgRNA sequences targeting PPP3CA or PPP3R1 were designed through VBC Score (vbc-score.org) and cloned into the PM04 plasmid, and sequentially inserted into pAAV-CMV-mScarlet-WPRE using Gibson assembly technology. sgRNA sequences were shown as follows:

PPP3CA-gRNA1: GACCATAGGATGTCACACAT;

PPP3CA-gRNA2: GCAGTCGAAGGCATCCATAC;

PPP3CA-gRNA3: GAGGCTGTTCGTACTTCTAC;

PPP3R1-gRNA1: GCTGATGAAATTAAAAGGCT;

PPP3R1-gRNA2: GCGATAAGGAACAGAAGTTG;

PPP3R1-gRNA3: GCAGAACCCTTTAGTACAGC.

### Viral injection

Mice at 8 weeks old were anaesthetized using 4-5% isoflurane (maintained at 1-2% for surgery), and 100uL virus (5×10^12^ gc/ml) was injected through retro-orbital sinus^111^. After two weeks to allow for expression, EEG implantation surgery was performed according to the protocol published previously ^51^. Mice were fixed in stereotaxic (RWD Life Science, 68405) and skull was exposed. Two holes were drilled at the frontal and the parietal cortex over the right cerebral hemisphere (Frontal: lateral to middle 1.5mm, 1.0mm anterior to bregma; parietal: lateral to middle 1.5 mm, 1.0 mm anterior to lambda). Two stainless steel screws (RWD Life Science), each soldered to a short copper wire, were inserted into the holes. Two EMG wires were implanted bilaterally into the neck muscle. All the copper wires were attached onto a micro-connector and was fixed to the skull. After surgery, mice were single housed for five days of recovery in new cages and then were placed into the special recording cage for three days to habituate to the recording cables.

### EEG and EMG recording and analysis

EEG and EMG data recording and analysis were performed as our previous study ^17^. EEG and EMG data at basal sleep conditions were recorded for 2 consecutive days, with a sample frequency of 200 Hz and epoch length of 4 seconds. EEG and EMG data were initially processed using AccuSleep ^112^ and then were manual correction in SleepSign. EEG and EMG signals were classified into Wake (fast and low amplitude EEG, high amplitude and variable EMG), NREM (high amplitude and 1-4 Hz dominant frequency EEG, low EMG tonus) and REM (low amplitude and 6-9 Hz frequency EEG, complete silent of EMG). The state episode was defined as at least three continuous and unitary state epochs. Epoch contained movement artifacts were included in sleep duration analysis but excluded from the subsequent power spectrum analysis. For power spectrum analysis, EEG was subjected to fast Fourier transform analysis (FFT). Power spectra represents the mean ratio of each 0.25 Hz to total 0–25 Hz of EEG signals during 24 hr baseline condition. The power density of NREMs represents the ratio of delta power density (1-4 Hz) to total power (0-25 Hz) in each hour. Cumulative rebound represented cumulative changes of time in post-SD compared with relative ZT under the baseline condition. Sleep/wake transition probabilities was analyzed as described in a previous study ^39^. For instance, *P_W to NR_* = N*_W to NR_* / (N*_W to W_* + N*_W to R_* + N*_W to NR_*), N*_W to NR_* denotes the number of transitions that transit from wakefulness epoch to NREM sleep epoch. W: wakefulness epoch, NR: NREM epoch, R: REM epoch.

### Sleep deprivation

After 2 consecutive days of EEG and EMG signals recording, mice were introduced into new cages at ZT0. Mice were gently handled or touched to keep them awake for 6 hrs of sleep deprivation, before being returned to the recording cage for another 24 hrs of recording.

### Statistical analysis

All statistical analyses were performed using GraphPad Prism 9.0. One-way ANOVA was used to compare differences among more than three groups, followed by Tukey’s multiple comparisons tests. Kruskal-Wallis tests were used for non-parameters tests. Two-way ANOVA was used to compare the differences between different groups with different treatments, followed by Tukey’s multiple comparisons tests. Two-way ANOVA with repeated measurements (Two-way RM ANOVA) was used when the same individuals were measured on the same outcome variable more than once, followed by Tukey’s multiple comparisons test. Data are presented as mean±SEM. In all cases, p values morethan 0.05 were considered not significant.

## Acknowledgements

We are grateful to Dr. Juan Huang at CIBR for generating SIK3 mice, Dr. Lei zhang at CIBR instrument core for help with customizing experimental devices for EEG recording, Dr. Yuan Li for help with mouse brain slice preparation, Linghao Kong and Ruixiang Wang from the CLS and Dr. Gongzheng Zhao from the Multi-Omics Mass Spectrometry Core of Shenzhen Bay Laboratory for assistance with mouse brain MS sample processing, the National Center for Protein Sciences at Peking University for access to instrumentation, and CLS, CIBR, CIMR, Changping Laboratory and the Chinese Academy of Medical Sciences (2019RU003) for support.

## Legends for Extended Data Figures and Tables

**Extended Data Fig. 1| Additional sleep phenotype of male T469A mutant mice. a-d,** Profiles of wake over 24 hrs (**a**), total wake duration over 24 hrs (**b**), wake episode number (**c**), wake episode duration (**d**) of SIK3^T469A/+^ (blue, n=10) and SIK3^+/+^ mice (black, n=11). *p<0.05; ns, not significant; mean±SEM (A: Two-way ANOVA with Tukey’s Multiple comparisons test; B: One-way ANOVA with Tukey’s multiple comparisons test; C-D: Kruskal-Wallis test with Dunn’s multiple comparisons test). **e**-**h,** Profiles of REM sleep over 24 hrs (**e**), total REM duration over 24 hrs (**f**), REM episode number (**g**), REM episode duration (**h**). *p<0.05; ns, not significant; mean ± SEM (E: Two-way ANOVA with Tukey’s multiple comparisons test; F: One-way ANOVA with Tukey’s multiple comparisons test; G-H: Kruskal-Wallis test with Dunn’s multiple comparisons test). **i**-**j,** EEG power spectrum during wake (**i**) or REM sleep (**j**). *p<0.05; mean±SEM (Two-way repeated measurement ANOVA with Tukey’s multiple comparisons test). **k**-**l,** Probabilities of transition between different sleep and wake states.

**Extended Data Fig. 2| Additional sleep phenotype of Male T469A mutant mice. a**-**b,** Recovery of wake (**a**) or REM sleep (**b**) after 6 hrs of SD. *p< 0.05; **p<0.01; ****p<0.0001; ns, not significant; mean ± SEM (Two-way ANOVA with Tukey’s multiple comparisons test). **c,** NREMS delta densities during the 24 hr recovery after SD. *p< 0.05; **p <0.01; ***p<0.001; ****p <0.0001; ns, not significant; mean ± SEM (Mixed-effects model). **d,** Changes of NREM delta power densities after SD. ns, not significant; mean ± SEM (Two-way repeated measurement ANOVA with Tukey’s multiple comparisons test). **e,** Representative hypnograms.

**Extended Data Fig. 3| Sleep phenotype of female T469A mutant mice.** Unlike males for which we could not obtain SIK3^T469A/T469A^ mutants, we did obtain all three genotypes for female mice: SIK3^T469A/T469A^ (red, n=5), SIK3^T469A/+^ (blue, n=6) and SIK3^+/+^ (black, n=7). **a-d,** Profiles of NREM over 24 hrs (**A**), total NREM duration over 24 hrs (**b**), NREM episode number (**c**), NREM episode duration (**d**). *p<0.05; ns, not significant; mean ± SEM (**a**: Two-way ANOVA with Tukey’s multiple comparisons test; **b**: One-way ANOVA with Tukey’s multiple comparisons test; **c**-**d**: Kruskal-Wallis test with Dunn’s multiple comparisons test). **e**-**h,** Profiles of REM sleep over 24 hrs (**e**), total REM duration over 24 hrs (**f**), REM episode number (**g**), REM episode duration (**h**). *p<0.05; **p <0.01; ns, not significant; mean ± SEM (**e**: Two-way ANOVA with Tukey’s multiple comparisons test; **f**: One-way ANOVA with Tukey’s multiple comparisons test; **g-h**: Kruskal-Wallis test with Dunn’s multiple comparisons test). **i-l,** Profiles of wake over 24 hrs (**i**), total wake duration over 24 hrs (**j**), wake episode number (**k**), wake episode duration (**l**). *p<0.05; ns, not significant; mean±SEM (**i**: Two-way ANOVA with Tukey’s multiple comparisons test; **j**: One-wayANOVA with Tukey’s multiple comparisons test; **k**-**l**: Kruskal-Wallis test with Dunn’s multiple comparisons test). **m**-**p,** Probabilities of transition between different sleep and wake states. *p<0.05; **p<0.01; ***p<0.001; ****p<0.0001; ns, not significant; mean±SEM (Two-way ANOVA with Tukey’s multiple comparisons test).

**Extended Data Fig. 4| Additional sleep phenotype of female T469A mutant mice. a**-**c,** EEG power spectrum during NREM sleep (**a**), REM sleep (**b**) or wake (**c**). *p< 0.05; mean ± SEM (Two-way repeated measurement ANOVA with Tukey’s multiple comparisons test). **d,** Diurnal NREM delta power densities. *p<0.05; **p<0.01; ***p<0.001; ****p<0.0001; ns, not significant; mean±SEM (Mixed-effects model). **e**-**g**, Recovery of NREM (**e**), REM (**f**) and wake (**g**) after 6 hrs of SD. ns, not significant; mean±SEM (Two-way ANOVA with Tukey’s multiple comparisons test). **h,** NREMS delta densities during the 24 hr recovery time. *p<0.05; **p<0.01; ***p<0.001; ****p <0.0001; ns, not significant; mean ± SEM (Mixed-effects model). **i,** Changes of NREM delta densities after 6 hrs of SD. *p< 0.05; **p<0.01; ns, not significant; mean±SEM (Two-way repeated measurement ANOVA with Tukey’s multiple comparisons test). **j,** One hr representative EEG and EMG signals of littermates at each vigilance state. **k,** Representative hypnograms.

**Extended Data Fig. 5| Specificities of antibodies against SIK3 T469 and SIK3 S551 phosphorylated by PKA.** Recombinant SIK3 fragment containing its amino acid residues 1 to 558 was expressed in and purified from *E. coli* before being treated by PKA in the presence of ATP. The PKA used was a mutant (PKA^T197E^) expressed in and purified from *E. coli*. Recombinant PKA^T197E^ phosphorylated SIK3 at T469 and S551, but not T221. Anti-phospho-SIK3^T469^ and anti-phospho-SIK3^S551^ antibodies specifically recognized T469 and S551, but not T221, of SIK3 under the same conditions.

**Extended Data Fig. 6| A schematic diagram of virally mediated gene knockdown in mice.** A host mouse either was WT or could express CAS9 from its Rosa26 site (Rosa26^Cas9/+^) and was injected with an AAV virus two weeks before an EEG recorder was placed on its head.

**Extended Data Fig. 7| Additional sleep phenotype of male PPP3CA knockdown mice. a**-**d,** Profiles of wake over 24 hrs (**a**), total wake duration over 24 hrs (**b**), wake episode number (**c**), wake episode duration (**d**). Wake was increased by approximately 3 hrs in PPP3CA^KD^ mice as compared to either control. Wake was increased during daytime by approximately 80 mins, due to increased wake episode duration (Extended Fig. 7d) albeit decreased wake episode number (Extended Data Fig. 7c) in PPP3CA^KD^ mice. Wake was significantly increased in nighttime by approximately 100 mins (Extended Data Fig. 7 a-b), due to increased duration (Extended Data Fig. 7d) albeit decreased number (Extended Data Fig. 7c) of wake episodes. *p<0.05; **p<0.01; ***p<0.001; ****p<0.0001; ns, not significant; mean±SEM (**a**: Two-way ANOVA with Tukey’s multiple comparisons test; **b**: One-way ANOVA with Tukey’s multiple comparisons test; **c-d**: Kruskal-Wallis test with Dunn’s multiple comparisons test. **e**-**h,** Profiles of REM sleep over 24 hrs (**e**), total REM duration over 24 hrs (**f**), REM episode number (**g**), REM episode duration (**h**). *p< 0.05; **p<0.01; ***p<0.001; ****p<0.0001; ns, not significant; mean±SEM (**e**: Two-way ANOVA with Tukey’s multiple comparisons test; **f**: One-way ANOVA with Tukey’s multiple comparisons test; **g**-**j**: Kruskal-Wallis test with Dunn’s multiple comparisons test). **i**-**j,** EEG power spectrum during wake and REM sleep. *p< 0.05; mean ± SEM (Two-way repeated measurement ANOVA with Tukey’s multiple comparisons test). **k**-**l,** Probabilities of transition between different sleep and wake states. *p< 0.05, **p<0.01, ***p<0.001 and ****p<0.0001, mean±SEM. (Two-way repeated measurement ANOVA with Tukey’s multiple comparisons test).

**Extended Data Fig. 8| Additional sleep phenotype of male PPP3CA knockdown mice. a**-**b,** Recovery of wake (**a**) and REM sleep (**b**) after 6 hrs of SD. *p< 0.05; **p <0.01; ***p<0.001; ****p<0.0001; ns, not significant; mean±SEM (Two-way ANOVA with Tukey’s multiple comparisons test). **c,** NREMS delta densities during the 24 hr recovery time. *p< 0.05; **p<0.01; ns, not significant; mean±SEM (Mixed-effects model). **d,** Changes of NREM delta densities after 6 hrs of SD. *p< 0.05; ns, not significant; mean±SEM (Two-way repeated measurement ANOVA with Tukey’s multiple comparisons test). **e,** Representative hypnograms.

**Extended Data Fig. 9| Additional sleep phenotype of PPP3R1 knockdown male mice. a-d,** Wake analysis: profiles showing wake time each hour in mins/hr with the X axis indicating ZT (**a**), total wake time over 24 hrs, wake time during the light phase (daytime) or the dark phase (**b**), wake episode number (**c**), wake episode duration (**d**). ns: statistically not significant; *p< 0.05; **p<0.01; ***p<0.001; ****p<0.0001; mean±SEM (a: Two-way ANOVA with Tukey’s multiple comparisons test; b: One-way ANOVA with Tukey’s multiple comparisons test; c-d: Kruskal-Wallis test with Dunn’s multiple comparisons test). **e-h,** REM analysis: profiles showing REM duration each hour in mins/hr with the X axis indicating ZT (**e**), total REM time over 24 hrs, REM time during the light phase (daytime) or the dark phase (**f**), REM episode number (**g**), REM episode duration (**h**). ns: statistically not significant; *p< 0.05; **p<0.01; ***p<0.001; ****p<0.0001; mean±SEM (**e**: Two-way ANOVA with Tukey’s multiple comparisons test; **f**: One-way ANOVA with Tukey’s multiple comparisons test; **g**-**h**: Kruskal-Wallis test with Dunn’s multiple comparisons test). (**i**-**j**) EEG power spectrum analysis of wake (**i**) and REM sleep (**j**). *p<0.05; mean±SEM (Two-way repeated measurement ANOVA with Tukey’s multiple comparisons test). **k**-**l,** Transition probabilities of different sleep and wake states. ns: statistically not significant; *p< 0.05; **p<0.01; ***p<0.001; ****p<0.0001; mean±SEM. (Two-way repeated measurement ANOVA with Tukey’s multiple comparisons test).

**Extended Data Fig. 10| Additional sleep phenotype of PPP3R1 knockdown male mice. a-b,** Recovery of umulative wake (**a**) and REMS b) after 6 hrs of SD. ns: statistically not significant; *p< 0.05; **p<0.01; ***p<0.001; ****p<0.0001; mean±SEM (Two-way ANOVA with Tukey’s multiple comparisons test). **c,** NREM delta power densities during the 24 hr recovery period after 6 hrs of SD. ns: statistically not significant; *p< 0.05; **p<0.01; ***p<0.001; ****p<0.0001;mean±SEM (Mixed-effects model). **d,** Changes of NREM delta power densities between pre- and post-SD. ns: statistically not significant; *p< 0.05; **p<0.01; ***p<0.001; ****p<0.0001; mean±SEM (Two-way repeated measurement ANOVA with Tukey’s multiple comparisons test). **e,** Representative hypnograms.

**Extended Data Table 1| Total Time spent in different sleep-wake states by SIK3^+/+^, SIK3^+/T469A^ and SIK3 ^T469A /T469A^ mice.**

**Extended Data Table 2| Differences in total time spent in different sleep-wake states by SIK3^+/+^, SIK3^+/T469A^ and SIK3 ^T469A/T469A^ Mice.**

**Extended Data Table 3| Total time spent in different sleep-wake States by eGFP^Ctrl^, WT^Ctrl^ and PPP3CA^KD^ mice.**

**Extended Data Table 4| Differences in total time spent in different sleep-wake states by eGFP^Ctrl^, WT^Ctrl^ and PPP3CA^KD^, ice.**

**Extended Data Table 5| Total time spent in different sleep-wake states by eGFP^Ctrl^, WT^Ctrl^ and PPP3R1^KD^ mice.**

**Extended Data Table 6| Differences in total time spent in different sleep-wake states by eGFP^Ctrl^, WT^Ctrl^ and PPP3R1^KD^ mice.**

