## Supplemental Table 1 for "Biochemical and chemical biological approaches to mammalian sleep: roles of calcineurin in site-specific dephosphorylation and sleep regulation"

### Table S1. Total time spent in different sleep-wake states of *Sik3*$^{+/+}$, *Sik3*$^{T469A/+}$ and *Sik3*$^{T469A/T469A}$ mice.

| | Total time | 24 hours (Mean ± SEM, min) | | Light Phase (Mean ± SEM, min) | | Dark Phase (Mean ± SEM, min) | |
|---|---|---|---|---|---|---|---|
| | | **Male** | **Female** | **Male** | **Female** | **Male** | **Female** |
| **Sleep** | *Sik3*$^{+/+}$ | 593.8 ± 14.2 | 503 ± 23.3 | 426.1 ± 5.5 | 392.2 ± 10.9 | 167.7 ± 11.2 | 110.3 ± 14.3 |
| | *Sik3*$^{T469A/+}$ | 643.9 ± 12.6 | 556.6 ± 32.3 | 433.8 ± 7.7 | 421.8 ± 7.4 | 210.1 ± 10.2 | 134.8 ± 28.5 |
| | *Sik3*$^{T469A/T469A}$ | | 578.8 ± 2.2 | | 377.5 ± 12.3 | | 201.3 ± 13.9 |
| | | **Male** | **Female** | **Male** | **Female** | **Male** | **Female** |
| **NREM** | *Sik3*$^{+/+}$ | 514.5 ± 12.8 | 427.2 ± 21.1 | 363.3 ± 4.4 | 328.6 ± 10.6 | 151.3 ± 10.5 | 98.7 ± 13.3 |
| | *Sik3*$^{T469A/+}$ | 570.3 ± 12.0 | 485.1 ± 30.9 | 378.3 ± 7.2 | 362.6 ± 6.8 | 192.0 ± 8.9 | 122.5 ± 26.4 |
| | *Sik3*$^{T469A/T469A}$ | | 520.1 ± 3.4 | | 336.9 ± 10.5 | | 183.2 ± 11.2 |
| | | **Male** | **Female** | **Male** | **Female** | **Male** | **Female** |
| **REM** | *Sik3*$^{+/+}$ | 79.3 ± 3.1 | 76.2 ± 4.1 | 62.9 ± 2.1 | 64.7 ± 3.1 | 16.4 ± 1.9 | 11.6 ± 1.1 |
| | *Sik3*$^{T469A/+}$ | 73.6 ± 2.1 | 71.6 ± 2.5 | 55.5 ± 1.7 | 59.3 ± 2.2 | 18.0 ± 1.9 | 12.3 ± 2.5 |
| | *Sik3*$^{T469A/T469A}$ | | 58.7 ± 2.5 | | 40.6 ± 2.7 | | 18.0 ± 3.1 |
| | | **Male** | **Female** | **Male** | **Female** | **Male** | **Female** |
| **Wake** | *Sik3*$^{+/+}$ | 846.2 ± 14.2 | 936.5 ± 23.3 | 293.9 ± 5.5 | 326.8 ± 10.9 | 552.3 ± 11.2 | 609.7 ± 14.3 |
| | *Sik3*$^{T469A/+}$ | 796.1 ± 12.6 | 883.5 ± 32.3 | 286.2 ± 7.7 | 298.2 ± 7.4 | 509.9 ± 10.2 | 585.3 ± 28.5 |
| | *Sik3*$^{T469A/T469A}$ | | 861.2 ± 2.2 | | 342.3 ± 12.3 | | 518.7 ± 13.9 |
