## Supplemental Table 2 for "Biochemical and chemical biological approaches to mammalian sleep: roles of calcineurin in site-specific dephosphorylation and sleep regulation"

**Table S2. Differences of total time spent in different sleep-wake states among *Sik3*$^{+/+}$, *Sik3*$^{T469A/+}$ and *Sik3*$^{T469A/T469A}$ mice.**

| | Differences | 24 hours (Mean ± SEM, min) | | Light Phase (Mean ± SEM, min) | | Dark Phase (Mean ± SEM, min) | |
|---|---|---|---|---|---|---|---|
| Sleep | | Male | Female | Male | Female | Male | Female |
| | *Sik3* $^{T469A/+}$ vs *Sik3* $^{+/+}$ | 50.1 ± 18.9 | 53.1 ± 33.4 | 7.7 ± 9.7 | 28.6 ± 14.1 | 42.4 ± 15.1 | 24.5 ± 27.6 |
| | *Sik3* $^{T469A/T469A}$ vs *Sik3* $^{+/+}$ | | 75.3 ± 35.2 | | -15.7 ± 14.9 | | 91.1 ± 29.0 |
| | *Sik3* $^{T469A/T469A}$ vs *Sik3* $^{T469A/+}$ | | 22.3 ± 36.4 | | -44.3 ± 15.4 | | 66.6 ± 30.0 |
| NREM | | Male | Female | Male | Female | Male | Female |
| | *Sik3* $^{T469A/+}$ vs *Sik3* $^{+/+}$ | 55.8 ± 17.5 | 57.8 ± 31.3 | 15.1 ± 8.7 | 34.0 ± 13.1 | 40.6 ± 13.6 | 23.8 ± 25.2 |
| | *Sik3* $^{T469A/T469A}$ vs *Sik3* $^{+/+}$ | | 92.9 ± 33.0 | | 8 ± 13.8 | | 84.6 ± 26.5 |
| | *Sik3* $^{T469A/T469A}$ vs *Sik3* $^{T469A/+}$ | | 35.1 ± 34.1 | | -25.7 ± 14.3 | | 60.8 ± 27.4 |
| REM | | Male | Female | Male | Female | Male | Female |
| | *Sik3* $^{T469A/+}$ vs *Sik3* $^{+/+}$ | -5.7 ± 3.7 | -4.7 ± 4.6 | -7.3 ± 2.7 | -5.4 ± 3.8 | 1.7 ± 2.7 | 0.7 ± 3.0 |
| | *Sik3* $^{T469A/T469A}$ vs *Sik3* $^{+/+}$ | | -17.6 ± 4.8 | | -24.0 ± 4.0 | | 6.5 ± 3.2 |
| | *Sik3* $^{T469A/T469A}$ vs *Sik3* $^{T469A/+}$ | | -12.9 ± 5.0 | | -18.6 ± 4.1 | | 5.7 ± 3.3 |
| Wake | | Male | Female | Male | Female | Male | Female |
| | *Sik3* $^{T469A/+}$ vs *Sik3* $^{+/+}$ | -50.1 ± 18.9 | -53.1 ± 33.4 | -7.7 ± 9.7 | -28.6 ± 14.1 | -42.4 ± 15.1 | -24.5 ± 27.6 |
| | *Sik3* $^{T469A/T469A}$ vs *Sik3* $^{+/+}$ | | -75.3 ± 35.2 | | 15.7 ± 14.86 | | -91.1 ± 29.0 |
| | *Sik3* $^{T469A/T469A}$ vs *Sik3* $^{T469A/+}$ | | -22.3 ± 36.4 | | 44.3 ± 15.4 | | -66.6 ± 30.0 |
