## Supplemental Table 3 for "Biochemical and chemical biological approaches to mammalian sleep: roles of calcineurin in site-specific dephosphorylation and sleep regulation"

### Table S3. Total time spent in different sleep-wake states of *eGFP*$^{Ctrl}$, *WT*$^{Ctrl}$ and *PPP3CA*$^{KD}$ mice

| | Total time | 24 hours (Mean ± SEM, min) | Light Phase (Mean ± SEM, min) | Dark Phase (Mean ± SEM, min) |
|---|---|---|---|---|
| **Sleep** | ***eGFP***$^{Ctrl}$ **mice** | 583.4 ± 9.0 | 428.9 ± 6.1 | 154.5 ± 7.6 |
| | ***WT***$^{Ctrl}$ **mice** | 588.4 ± 16.0 | 433.6 ± 7.4 | 154.8 ± 12.7 |
| | ***PPP3CA***$^{KD}$ **mice** | 395.8 ± 7.0 | 349.4 ± 4.6 | 46.4 ± 4.9 |
| **NREM** | ***eGFP***$^{Ctrl}$ **mice** | 504.6 ± 8.6 | 364.6 ± 6.1 | 139.9 ± 7.0 |
| | ***WT***$^{Ctrl}$ **mice** | 510.5 ± 15.6 | 368.8 ± 7.6 | 141.7 ± 11.5 |
| | ***PPP3CA***$^{KD}$ **mice** | 303.5 ± 6.3 | 263.8 ± 4.3 | 39.7 ± 3.9 |
| **REM** | ***eGFP***$^{Ctrl}$ **mice** | 78.9 ± 1.8 | 64.3 ± 1.6 | 14.6 ± 1.2 |
| | ***WT***$^{Ctrl}$ **mice** | 77.9 ± 2.2 | 64.9 ± 2.2 | 13.0 ± 1.3 |
| | ***PPP3CA***$^{KD}$ **mice** | 92.2 ± 1.9 | 85.6 ± 1.7 | 6.7 ± 1.1 |
| **Wake** | ***eGFP***$^{Ctrl}$ **mice** | 856.6 ± 9.0 | 291.1 ± 6.1 | 565.5 ± 7.6 |
| | ***WT***$^{Ctrl}$ **mice** | 851.6 ± 16.0 | 286.4 ± 7.4 | 565.2 ± 12.7 |
| | ***PPP3CA***$^{KD}$ **mice** | 1044.2 ± 7.0 | 370.6 ± 4.6 | 673.6 ± 4.9 |
