## Supplemental Table 4 for "Biochemical and chemical biological approaches to mammalian sleep: roles of calcineurin in site-specific dephosphorylation and sleep regulation"

### Table S4. Differences of total time spent in different sleep-wake states among *eGFP*$^{Ctrl}$, *WT*$^{Ctrl}$ and *PPP3CA*$^{KD}$ mice

| | Differences | 24 hours (Mean ± SEM, min) | Light Phase (Mean ± SEM, min) | Dark Phase (Mean ± SEM, min) |
|---|---|---|---|---|
| **Sleep** | ***PPP3CA***$^{KD}$ ***vs eGFP***$^{Ctrl}$ | -187.6 ± 13.0 | -79.5 ± 7.7 | -108.1 ± 10.3 |
| | ***PPP3CA***$^{KD}$ ***vs WT***$^{Ctrl}$ | -192.6 ± 14.7 | -84.2 ± 8.7 | -108.4 ± 11.6 |
| | ***eGFP***$^{Ctrl}$ ***vs WT***$^{Ctrl}$ | 5.0 ± 15.1 | 4.8 ± 8.9 | 0.2 ± 11.9 |
| **NREM** | ***PPP3CA***$^{KD}$ ***vs eGFP***$^{Ctrl}$ | -201.0 ± 12.4 | -100.8 ± 7.6 | -100.2 ± 9.2 |
| | ***PPP3CA***$^{KD}$ ***vs WT***$^{Ctrl}$ | -207.0 ± 14.0 | -104.9 ± 8.5 | -102.0 ± 10.4 |
| | ***eGFP***$^{Ctrl}$ ***vs WT***$^{Ctrl}$ | -5.9 ± 14.4 | -4.1 ± 8.8 | -1.8 ± 10.7 |
| **REM** | ***PPP3CA***$^{KD}$ ***vs eGFP***$^{Ctrl}$ | 13.3 ± 2.6 | 21.3 ± 2.3 | -7.9 ± 1.6 |
| | ***PPP3CA***$^{KD}$ ***vs WT***$^{Ctrl}$ | 14.4 ± 2.9 | 20.7 ± 2.6 | -6.3 ± 1.8 |
| | ***eGFP***$^{Ctrl}$ ***vs WT***$^{Ctrl}$ | 1.0 ± 3.0 | -0.5 ± 2.7 | 1.6 ± 1.8 |
| **Wake** | ***PPP3CA***$^{KD}$ ***vs eGFP***$^{Ctrl}$ | 187.6 ± 13.0 | 79.5 ± 7.7 | 108.1 ± 10.3 |
| | ***PPP3CA***$^{KD}$ ***vs WT***$^{Ctrl}$ | 192.6 ± 14.7 | 84.2 ± 8.7 | 108.4 ± 11.6 |
| | ***eGFP***$^{Ctrl}$ ***vs WT***$^{Ctrl}$ | -5.0 ± 15.1 | -4.8 ± 8.9 | -0.2 ± 11.9 |
