## Supplemental Table 5 for "Biochemical and chemical biological approaches to mammalian sleep: roles of calcineurin in site-specific dephosphorylation and sleep regulation"

**Table S5. Total time spent in different sleep-wake states of *eGFP*$^{Ctrl}$, *WT*$^{Ctrl}$, and *PPP3R1*$^{KD}$ mice**

| | Total time | 24 hours (Mean ± SEM, min) | Light phase (Mean ± SEM, min) | Dark phase (Mean ± SEM, min) |
|---|---|---|---|---|
| **Sleep** | ***eGFP*$^{Ctrl}$ mice** | 617.8 ± 6.6 | 438.5 ± 5.3 | 179.4 ± 10.0 |
| | ***WT*$^{Ctrl}$ mice** | 585.7 ± 11.3 | 441.4 ± 5.1 | 144.3 ± 9.7 |
| | ***PPP3R1*$^{KD}$ mice** | 236.4 ± 24.5 | 215.9 ± 24.3 | 20.4 ± 8.5 |
| **NREM** | ***eGFP*$^{Ctrl}$ mice** | 543.6 ± 6.5 | 378.0 ± 4.6 | 165.7 ± 8.9 |
| | ***WT*$^{Ctrl}$ mice** | 512.0 ± 10.6 | 378.5 ± 4.0 | 133.5 ± 9.5 |
| | ***PPP3R1*$^{KD}$ mice** | 183.6 ± 18.3 | 165.1 ± 17.3 | 18.47 ± 7.7 |
| **REM** | ***eGFP*$^{Ctrl}$ mice** | 74.2 ± 1.5 | 60.6 ± 1.0 | 13.6 ± 1.8 |
| | ***WT*$^{Ctrl}$ mice** | 73.8 ± 2.9 | 62.9 ± 2.6 | 10.9 ± 1.2 |
| | ***PPP3R1*$^{KD}$ mice** | 52.8 ± 8.3 | 50.9 ± 8.5 | 1.9 ± 0.9 |
| **WAKE** | ***eGFP*$^{Ctrl}$ mice** | 822.2 ± 6.6 | 281.5 ± 5.3 | 540.7 ± 10.0 |
| | ***WT*$^{Ctrl}$ mice** | 854.3 ± 11.3 | 278.6 ± 5.1 | 575.6 ± 9.7 |
| | ***PPP3R1*$^{KD}$ mice** | 1204.0 ± 24.5 | 504.0 ± 24.3 | 699.6 ± 8.5 |
