## Supplemental Table 6 for "Biochemical and chemical biological approaches to mammalian sleep: roles of calcineurin in site-specific dephosphorylation and sleep regulation"

**Table S6. Differences of total time spent in different sleep-wake states among *eGFP*$^{Ctrl}$, *WT*$^{Ctrl}$, and *PPP3R1*$^{KD}$ mice**

| | Differences | 24 hours (Mean ± SEM, min) | Light phase (Mean ± SEM, min) | Dark phase (Mean ± SEM, min) |
|---|---|---|---|---|
| **Sleep** | ***PPP3R1*$^{KD}$ *vs eGFP*$^{Ctrl}$** | -381.4 ± 22.8 | -222.6 ± 20.0 | -159.0 ± 14.1 |
| | ***PPP3R1*$^{KD}$ *vs WT*$^{Ctrl}$** | -349.3 ± 21.5 | -225.5 ± 18.9 | -123.8 ± 13.3 |
| | ***eGFP*$^{Ctrl}$ *vs WT*$^{Ctrl}$** | 32.1 ± 21.5 | -2.9 ± 18.9 | 35.1 ± 13.3 |
| **NREM** | ***PPP3R1*$^{KD}$ *vs eGFP*$^{Ctrl}$** | -360.1 ± 18.5 | -212.9 ± 14.5 | -147.2 ± 13.2 |
| | ***PPP3R1*$^{KD}$ *vs WT*$^{Ctrl}$** | -328.4 ± 17.5 | -213.4 ± 13.7 | -115.0 ± 12.5 |
| | ***eGFP*$^{Ctrl}$ *vs WT*$^{Ctrl}$** | 31.7 ± 17.5 | -0.6 ± 13.7 | 32.2 ± 12.5 |
| **REM** | ***PPP3R1*$^{KD}$ *vs eGFP*$^{Ctrl}$** | -21.4 ± 7.2 | -9.1 ± 7.1 | -11.7 ± 2.0 |
| | ***PPP3R1*$^{KD}$ *vs WT*$^{Ctrl}$** | -20.9 ± 6.8 | -12.0 ± 6.7 | -9.0 ± 1.8 |
| | ***eGFP*$^{Ctrl}$ *vs WT*$^{Ctrl}$** | 0.4 ± 6.8 | -2.3 ± 6.7 | 2.8 ± 1.8 |
| **WAKE** | ***PPP3R1*$^{KD}$ *vs eGFP*$^{Ctrl}$** | 381.4 ± 22.8 | 222.6 ± 20.1 | 158.9 ± 14.1 |
| | ***PPP3R1*$^{KD}$ *vs WT*$^{Ctrl}$** | 349.3 ± 21.5 | 225.4 ± 19.0 | 123.9 ± 13.3 |
| | ***eGFP*$^{Ctrl}$ *vs WT*$^{Ctrl}$** | -32.1 ± 21.5 | 2.9 ± 19.0 | -34.9 ± 13.3 |
